# Reprogramming BCMA-Targeted CAR T Cells through γ-Secretase Modulation Blocks Antigen Shedding and Extends CAR T Longevity

**DOI:** 10.64898/2026.01.20.700575

**Authors:** Sheetal Sharma, Varnit Chauhan, Zahoor Ahmad Bhat, Jyotirmoi Aich, Divya, Sunita Gupta, Divya Rashmi, Ruquaiya Alam, Mohammad Sufyan Ansari, Drashti Dave, Preet Patel, Nisha Chaudhary, Deepak Kumar Singh, Vijay Pal Singh, Mohammad Husain, Amit Srivastava Kumar, Ulaganathan Mabalirajan, Dinesh Pendharkar, Vishnu Hari, Nedunchezhian Murugaiyan, Soumya Sinha Roy, Sivaprakash Ramalingam, Gaurav Kharya, Amit Kumar Verma, Tanveer Ahmad

## Abstract

B-cell maturation antigen (BCMA) shedding by γ-secretase generates soluble BCMA (sBCMA), which diminishes membrane BCMA (mBCMA) antigen density, and limits the durability of BCMA-directed immunotherapies in multiple myeloma (MM). Here, using AI-driven diffusion modeling, we identify peptide inhibitors that selectively block γ-secretase-mediated BCMA cleavage, stabilizing mBCMA without compromising cellular viability. The lead peptide, P5, suppresses sBCMA and restores mBCMA in vitro and in xenograft models. To create sustained, cell-intrinsic inhibition, we engineered Shedding Deterrent Locked-in (SHEDLOCK) CAR T cells that locally secrete γ-secretase-modulatory peptides. SHEDLOCK-1 CAR T cells secreting P5 exhibit enhanced cytotoxicity and persistence relative to conventional CAR T cells. Building on this, SHEDLOCK-2 CAR T cells were generated, which secrete a modified, naturally derived peptide (nxP) that modulates γ-secretase activity, concurrently preventing BCMA shedding, and surprisingly enhancing CAR T cell longevity by preserving telomere integrity and metabolic fitness. In MM xenograft and patient-derived models, SHEDLOCK-2 CAR T cells demonstrate durable antitumor activity with a favorable safety profile. Together, these findings establish SHEDLOCK as a next-generation CAR T platform and provide a strong preclinical foundation for Phase I clinical evaluation.

## Introduction

BCMA has emerged as one of the most reliable and selective therapeutic targets in MM because it is largely restricted to malignant plasma cells and plays a key role in sustaining their growth through NF-κB and PI3K-AKT signaling^1^. Although BCMA directed therapies including CAR T cells, bispecific antibodies, newer T cell engagers and antibody drug conjugates have shown impressive clinical activity, relapse over time is still common and remains a major challenge^2, 3^. Clinical outcomes from trials of the two FDA approved therapies; idecabtagene vicleucel (ide-cel; bb2121) and ciltacabtagene autoleucel (cilta-cel) demonstrate overall response rates exceeding 80%; however, the median progression-free survival rarely surpasses 12 months, suggesting the limited persistence and functional decline of BCMA-directed CAR T cells with time^4–6^ . Thus, understanding and overcoming the biological mechanisms related to this loss of efficacy, particularly BCMA shedding, antigen escape, and T cell senescence, have remained critical unmet challenges.

BCMA exists as a type III transmembrane glycoprotein that binds the ligands APRIL and BAFF, transmitting pro-survival signals through NF-κB, AKT, and MAPK pathways^7, 8^. However, this receptor undergoes proteolytic cleavage by γ-secretase, which cleaves its extracellular domain, releasing sBCMA into circulation^9, 10^. The consequence is twofold: (1) a reduction of mBCMA density on myeloma cells, diminishing CAR T cell engagement and immune synapse formation, and (2) the generation of circulating sBCMA, which acts as a decoy receptor capable of binding and neutralizing anti-BCMA CARs or bispecific antibodies^10–12^. Clinically, higher circulating sBCMA concentrations exceeding in MM patients have been observed and shown to correlate with lower antigen density and reduced efficacy of BCMA-targeting therapies^10, 13^. In a prospective study of relapsed/refractory MM patients treated with ide-cel, sBCMA levels dropped sharply within 10 days of infusion in responders but rebounded upon CAR T contraction, marking disease recurrence^14^ . Mechanistically, presence of high levels of sBCMA functionally impairs immune therapy by blocking CAR recognition sites. Multiple studies have provided compelling biochemical evidence that high circulating sBCMA competitively inhibits BCMA-targeting bispecific antibody binding, reducing immune synapse formation and T cell activation^12^ . In parallel, Lee et al. (2024)^15^ and Chen et al. (2022)^12^ demonstrated that sBCMA acts as a key determinant of therapeutic refractoriness to BCMA T cell engagers by sequestering immune effectors and altering cytokine dynamics.

To overcome these challenges, previous studies have used a rational approach of inhibiting γ-secretase to prevent BCMA cleavage and established that BCMA shedding is directly dependent on γ-secretase enzymatic activity^11^. Further, translational studies demonstrated that pharmacologic γ-secretase inhibitors (GSIs) dramatically increase mBCMA surface levels and enhance sensitivity to BCMA CAR T cells and bispecific antibodies^10, 12, 16^. Notably, preclinical studies combining GSIs with CAR T cells have shown elevated BCMA surface expression up to 150-fold while decreasing sBCMA, leading to augmented CAR T activation and cytokine release^10, 12^. Early-phase clinical trials (NCT03502577) combining GSI with BCMA CAR T cell therapy confirmed these findings, reporting improved expansion and tumor clearance without significantly increasing toxicity^16–18^. The synergistic augmentation of CAR T cell function through γ-secretase blockade thus represents one of the first clinically validated strategies to counteract antigen escape and enhance therapeutic durability. Further, the clinical correlation of sBCMA-mediated resistance has become increasingly clear. Patients with persistently elevated sBCMA following ide-cel therapy exhibited reduced CAR expansion and earlier relapse, with plasma sBCMA elevating concurrently with disease progression^14^. Importantly, both sBCMA level and shedding kinetics have been validated as dynamic biomarkers predicting relapse and loss of response with time^19^. Together, these studies establish a clear mechanistic link between proteolytic BCMA loss and clinical refractoriness.

The other challenge beside BCMA antigen loss, is that even in patients with adequate antigen expression, the persistence of BCMA CAR T cells remains limited. Longitudinal immune monitoring of patients treated with ide-cel or cilta-cel reveals rapid expansion followed by contraction to near-undetectable levels within months^5, 20^. This short-lived persistence correlates with loss of central memory subsets, mitochondrial dysfunction, and progressive metabolic exhaustion^21, 22^. Transcriptomic analyses of post-infusion CAR T cells have identified upregulation of senescence-associated markers such as CDKN1A (p21) and CDKN2A (p16), along with increased reactive oxygen species and shortened telomere length^23^. Functionally, these changes drive loss of proliferative capacity and cytokine secretion, which are hallmarks of immune aging within the tumor milieu^24^. Recent studies have demonstrated the role of metabolic programming and Notch signaling in shaping T cell fate^25, 26^. Sustained Notch activation in T cells accelerates differentiation towards terminal effector phenotypes and suppresses telomerase activity, predisposing them to senescence and apoptosis^27, 28^. Conversely, metabolic interventions that restore mitochondrial biogenesis, regulate mTORC1 activity or inhibit aberrant Notch signaling, can promote memory-like persistence and functional rejuvenation^29–33^. The integration of metabolic and signaling modulation thus represents an emerging paradigm for next-generation CAR design.

To extend the functional lifespan of CAR T cells, multiple strategies have been explored: (1) genetic engineering of CAR constructs to resist tonic signaling and activation-induced exhaustion, (2) metabolic reprogramming via PGC1α activation or AKT3 inhibition to sustain mitochondrial health, and (3) co-administration of GSIs or soluble decoy traps to maintain antigen availability^34–37^. However, no single approach simultaneously addresses both antigen stability and T cell longevity.

Addressing these limitations, we developed cell-intrinsic, peptide-based γ-secretase inhibitors that block BCMA shedding and enhance CAR T cell metabolic fitness. Incorporating these peptides into CAR T cells improved surface BCMA retention, reduced sBCMA decoy interference, and sustained cytotoxicity while mitigating senescence and increasing longevity via Notch signaling modulation. This approach mechanistically links BCMA shedding with T cell aging and exhaustion, as well as metabolic decline, providing a unified strategy to achieve durable BCMA CAR T cell efficacy.

## Results

### De novo designed peptides inhibit γ-secretase mediated BCMA shedding

Using target-specific generative artificial intelligence based on RFdiffusion, we designed a library of candidate sequences predicted to bind the γ-secretase cleavage site within the transmembrane (TM) region of BCMA (**Figure 1A** and **Table 1**). Initial computational screening identified five top-ranking peptide binders (P01 to P05) that were chosen for their high structural confidence and strong complementarity to the BCMA TM region (**Figure 1B**). The peptide generation workflow is described in the Methods. We next used structural modeling and all-atom molecular dynamics (MD) simulations to examine how each peptide engaged the BCMA TM helix, the region that γ-secretase cleaves^11, 38, 39^. Over a 200 ns simulation, every peptide held stable contacts with the TM helix through hydrophobic packing and occasional hydrogen bonds (**Figure 1C and 1D, Supplementary Figure S1A, and Movie 1**). Replicate simulations showed that the peptides consistently remained beside the TM helix within a POPC:DOPC lipid bilayer for most of the simulated time (**Figure 1E and 1F, and Supplementary Figure S1B to S1F**). Per-residue energy decomposition of each peptide-BCMA complex is shown in **Supplementary Figure S2**.

**Figure 1:**
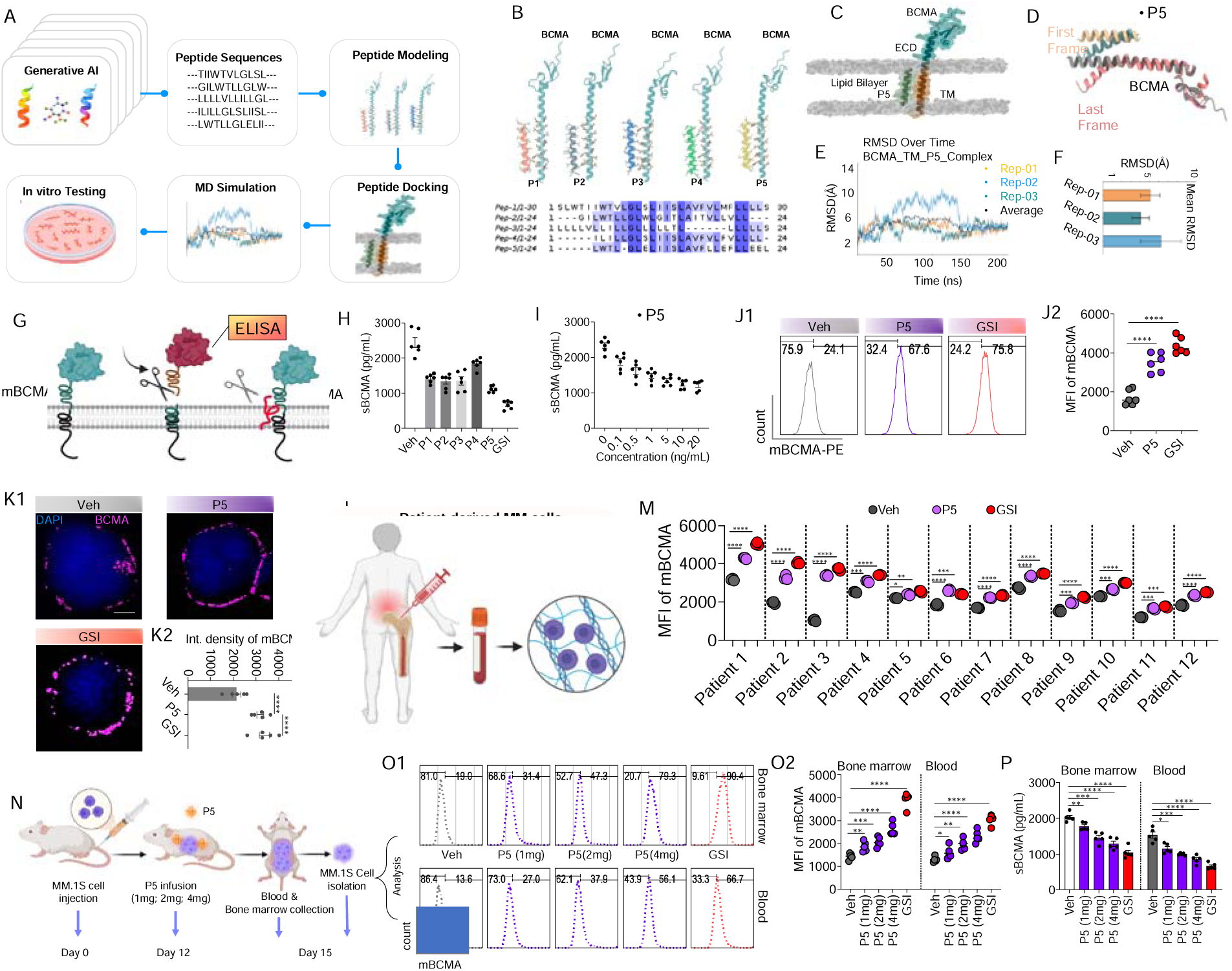
De novo peptides inhibit γ-secretase-mediated BCMA shedding. (A) Workflow for peptide generative modeling and in-silico screening while targeting the γ-secretase cleavage region of BCMA transmembrane domain. (B) Structural representation and sequences of the shortlisted peptides; P1-P5 from computational evaluation. (C-D) Structural models showing P5 interacting with the BCMA transmembrane helix within a POPC:DOPC lipid bilayer. (E) Data from MD simulations wherein root mean square deviation (RMSD) shows the evolution of BCMA-P5 complex over course of 200 ns revealing the structural convergence of the complexes. (F) The average RMSD of three replicates. (G) In vitro BCMA shedding experimental model. (H-I) Soluble BCMA ELISA quantification with the different in-silico generated peptides and dose-dependent reduction by best performing peptide, P5 in functional assay. (J1) Representative flow cytometry histogram plots showing sustained membrane BCMA (mBCMA) retention after P5 treatment; GSI DAPT serves as the positive control. (J2) Corresponding bar graph analysis of the flow cytometry data (n=6). (K1) Super-resolution microscopy images in P5-treated MM.1S cells showing retention of mBCMA (magenta) as shown by retained surface expression. DAPI (blue) was used to label nuclei. (K2) Bar graphs of the images showing integrated mBCMA density for the expression shown in K1 (n=5). (L) Workflow of isolation and culture of primary multiple myeloma (MM) cells from patient bone marrow. (M) Flow cytometry analysis of patient-derived MM samples showing increased mBCMA after P5 treatment (n=12 patient samples); the corresponding sBCMA ELISA is shown in Supplementary Figure S5. (N) NSG MM xenograft study design for systemic P5 testing. (O1) Representative histograms showing the in vivo increase of tumor-cell mBCMA in bone marrow and blood after P5 treatment. (O2) Corresponding quantification of mBCMA (n=5). (P) ELISA of sBCMA in bone marrow and blood showing dose-dependent reduction after P5 treatment. Data represent mean ± SEM. *p < 0.05; **p < 0.01; ***p < 0.001; ****p < 0.0001. A two-tailed Mann–Whitney U test was used for statistical analysis between groups.

To evaluate this functionally, we established an in vitro BCMA shedding assay and used GSI DAPT as a control (**Figure 1G**). sBCMA measured by ELISA showed that P05 (P5 hereafter) lowered sBCMA release in a dose-dependent manner more effectively than the other variants (**Figure 1H and 1I**). Flow cytometry confirmed that P5 preserved mBCMA on the cell surface (**Figure 1J1 and 1J2),** and super-resolution microscopy imaging revealed a dense, organized array of surface BCMA receptors in treated cells **(Figure 1K1 and 1K2).** In BCMA-stable HEK293 and K562 cells (HEK-BCMA and K562-BCMA), P5 raised mBCMA and reduced sBCMA (**Supplementary Figure S3**). P5 did not affect proliferation or viability in epithelial, B cell, T cell, NK cell, and neuronal lineages, indicating specificity and no off-target toxicity (**Supplementary Figure S4**).

To assess the clinical relevance of P5, MM cells were isolated from 12 patient bone marrow biopsy samples, enriched with the RosetteSep Human MM Cell Enrichment Kit and cultured with P5 (**Figure 1L**). In all twelve samples, P5 promoted retention of mBCMA and lowered sBCMA in the culture supernatant (**Figure 1M** and **Supplementary Figure S5**).

We then tested P5 in vivo using an NSG xenograft mice model of MM established with MM.1S cells (**Figure 1N**). Systemic infusion of P5 produced a dose-dependent increase in mBCMA on MM.1S cells (**Figure 1O1 and 1O2**) and a dose-dependent fall in circulating and bone marrow sBCMA (**Figure 1P**). Because these effects appeared in both peripheral blood and bone marrow, the response to P5 was broad and sustained. Together, these findings show that the de novo designed peptide P5 selectively targets the BCMA TM region and suppresses γ-secretase mediated cleavage. By stabilizing mBCMA and limiting sBCMA shedding, P5 supports lasting receptor retention, and we therefore prioritized it for further study.

### Humanized CAR T cells show improved binding kinetics and potent anti-tumor activity

The single-chain variable fragment (scFv) strongly influences CAR T cell efficacy, yet murine scFvs tend to provoke human anti-mouse antibody (HAMA) responses that can shorten therapeutic durability^40, 41^. To address this, we applied an AI-guided design pipeline to generate humanized scFvs that lower immunogenicity while keeping high-affinity recognition of BCMA, including the low-density BCMA often seen in MM. Comparing the murine clone 4C8A with the AI-humanized variants showed a clear reduction in non-human sequence content alongside preserved CDRs. Per-residue energy decomposition of the murine and humanized BCMA complexes showed that the CDR residues driving BCMA engagement were retained after humanization (**Supplementary Figure S6A and S6B**). Structural modeling showed that the humanized scFv kept the CDR loops correctly oriented toward the BCMA epitopes (**Figure 2A**), with stable complexes and comparable binding free energies across replicate simulations (**Figure 2B and 2C**), confirming that the substituted framework residues remained compatible with the original CDRs. Free energy perturbation (FEP) analysis quantified the affinity changes from point mutations within the CDRs and confirmed that epitope specificity and binding affinity were retained **(Supplementary Figure S6C and S6D),** and the hydrogen-bonding patterns of the substituted residues are shown in (**Supplementary Figure S6E to S6J)**. These in silico analyses establish AI-guided humanization, refined through repeated computational evaluation, as an effective way to engineer scFvs with better humanness while maintaining binding affinity and cytotoxicity.

**Figure 2:**
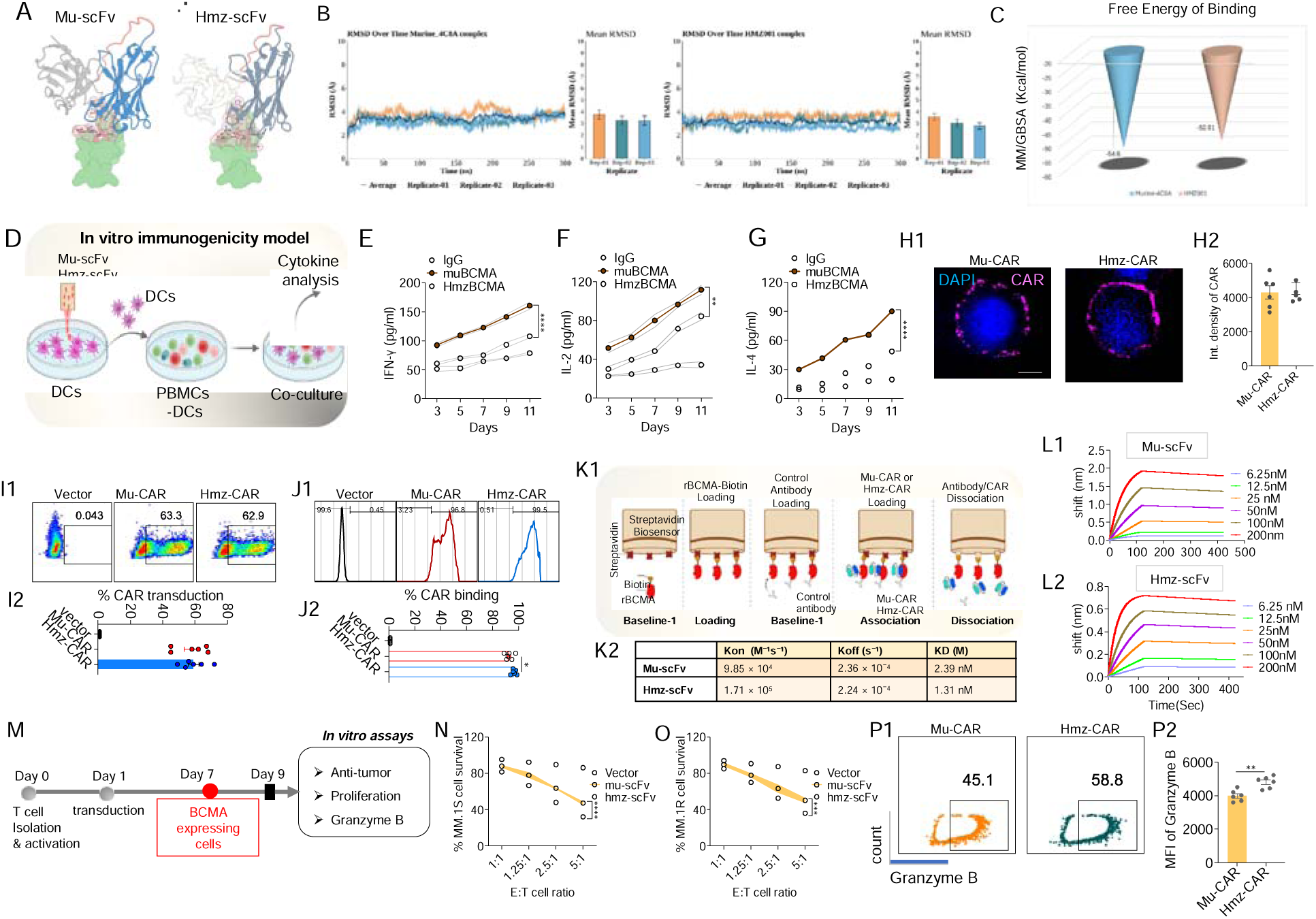
Humanized anti-BCMA CARs show reduced immunogenicity, improved kinetics, and enhanced cytotoxicity. (A) Structural modeling of murine 4C8A clone versus LLM-based humanized scFv variant showing preserved CDR orientation towards BCMA and specificity for the distinct epitope. (B) RMSD plots for the snapshots from MD simulation of the scFvs (murine and humanized) in complex with BCMA across three independent replicates. (C) MM/GBSA based free energy binding plots for the murine and humanized scFv-BCMA complexes. (D) Workflow of immunogenicity assay showing antibody-primed dendritic cells co-cultured with autologous PBMCs. (E-G) Cytokine analysis of IFN-γ, IL-2, and IL-4 showing lower levels for humanized CARs versus murine. Fully human IgG antibody was used as reference control. (H1) Representative super-resolution microscopy images showing uniform membrane localization of CAR constructs (magenta). Nuclei were stained with DAPI (blue). (H2) Corresponding bar graph of the image analysis (n=6). (I1) Dot plots of flow cytometry quantification of CAR surface expression. (I2) Bar graph of the flow cytometry data showing percentage CAR Transduction (n=6). (J1) Cell-based binding assay using flow cytometry showing affinity gain for the CDR-optimized humanized CAR (HmzCAR). (J2) Bar graph of the analysis (n=6). (K1) Workflow of the biolayer interferometry (BLI) assay. (K2) Binding kinetic constants (kon, koff and KD) for the murine and humanized scFv. (L1, L2) BLI sensograms showing real-time binding kinetics of the indicated analytes. Colored traces represent different concentrations, with an initial association phase followed by maintained dissociation. (M) Experimental workflow of co-culture of CAR T cells with BCMA expressing target cells. (N) Cytotoxicity against MM.1S cells across different E:T ratios (n=5). (O) Similarly, cytotoxicity against MM.1R cells at different E:T ratios (n=5). (P1) Representative flow cytometry contour plots of Granzyme B secretion. (P2) Mean fluorescence intensity (MFI) of the flow cytometry contour plots (n=6). Data represent mean ± SEM. *p < 0.05; **p < 0.01; ***p < 0.001; ****p < 0.0001. A two-tailed Mann–Whitney U test was used for statistical analysis between groups.

To gauge immunogenic potential, we co-cultured antibody-primed dendritic cells with autologous PBMCs (**Figure 2D**) and measured cytokine induction by CAR constructs carrying each scFv after antigen presentation. Humanized CARs triggered much lower IFN-γ, IL-2, and IL-4 secretion than the murine scFv, while a fully human IgG served as a low-risk reference (**Figure 2E to 2G**). These data confirm that humanization reduces T cell alloreactivity and lowers the risk of HAMA responses.

For functional testing, we generated CAR T cells as described previously ^42^. Super-resolution microscopy showed uniform membrane localization of CAR molecules across murine and humanized CAR T constructs **(Figure 2H1 and 2H2),** which was also confirmed by flow cytometry using a BCMA CAR detection antibody **(Figure 2I1 and 2I2)**. Binding assays indicated a modest affinity gain in the CDR-optimized humanized CAR, termed HmzCAR **(Figure 2J1 and 2J2).** Bio-layer interferometry (BLI) agreed, showing that HmzCAR bound with an affinity comparable to or slightly better than the murine scFv (KD 1.31 nM versus 2.39 nM **(Figure 2K1, 2K2, 2L1 and 2L2).**

We next measured cytotoxicity across several effector-to-target (E:T) ratios using MM.1S and MM.1R cells. HmzCAR T cells lysed tumor cells far more effectively than murine CARs (**Figure 2M to 2O**), in line with higher Granzyme B levels **(Figure 2P1 and 2P2).** In engineered HEK-BCMA and K562-BCMA models expressing different BCMA densities, HmzCAR T cells cleared both high- and low-BCMA density cell line models, whereas the murine CAR T cells were weaker against low-BCMA targets **(Supplementary Figure S7A, S7B1 and S7B2).** These results show that computationally designed HmzCAR T cells combine reduced immunogenicity, consistent binding affinity, and stronger anti-tumor potency.

### SHEDLOCK-1 CAR T cells enhance anti-myeloma efficacy by blocking BCMA shedding

To assess how sBCMA affects CAR T cell activity, we measured tumor cytotoxicity and cytokine secretion after exposing CAR T cells to supernatant from 12 independent patient-derived MM cell cultures (**Figure 3A**). Supernatants collected 24 h after MM culture were mixed 1:1 with CAR T cells, which were then co-cultured with MM.1S cells. CAR T cells exposed to MM supernatant showed a clear loss of cytolytic activity and of Granzyme B and IL-2 secretion, indicating that sBCMA in the medium inhibits CAR T function **(Figure 3B to 3D).** To counter this inhibition, we combined the BCMA-blocking peptide P5 with the HmzCAR construct. CAR T cells treated with rising concentrations of P5 gained function in a dose-dependent manner, and this gain was tracked with improved tumor killing (**Figure 3E and 3F**). To make the system self-contained, we engineered CAR T cells to secrete either a non-targeting peptide (NTP) or the P5 peptide as a built-in module, a strategy we named SHEDLOCK-1 (Shedding Deterrent Locked-in-1) CAR T cells (**Figure 3G**). We confirmed peptide expression and secretion by ELISA assay **(*methods*; Supplementary Figure S7C, D**) and by testing whether SHEDLOCK-1 supernatant blocked BCMA shedding. Flow cytometry showed increased mBCMA on MM cells treated with supernatant from SHEDLOCK-1 CAR T cells, confirming reduced BCMA shedding **(Figure 3H1 and 3H2),** and super-resolution imaging of MM.1S cells incubated with SHEDLOCK-1 CAR T cells supernatant showed strong mBCMA retention relative to controls **(Figure 3I1 and 3I2)**, verifying efficient peptide expression and activity in the SHEDLOCK-1 setting.

**Figure 3:**
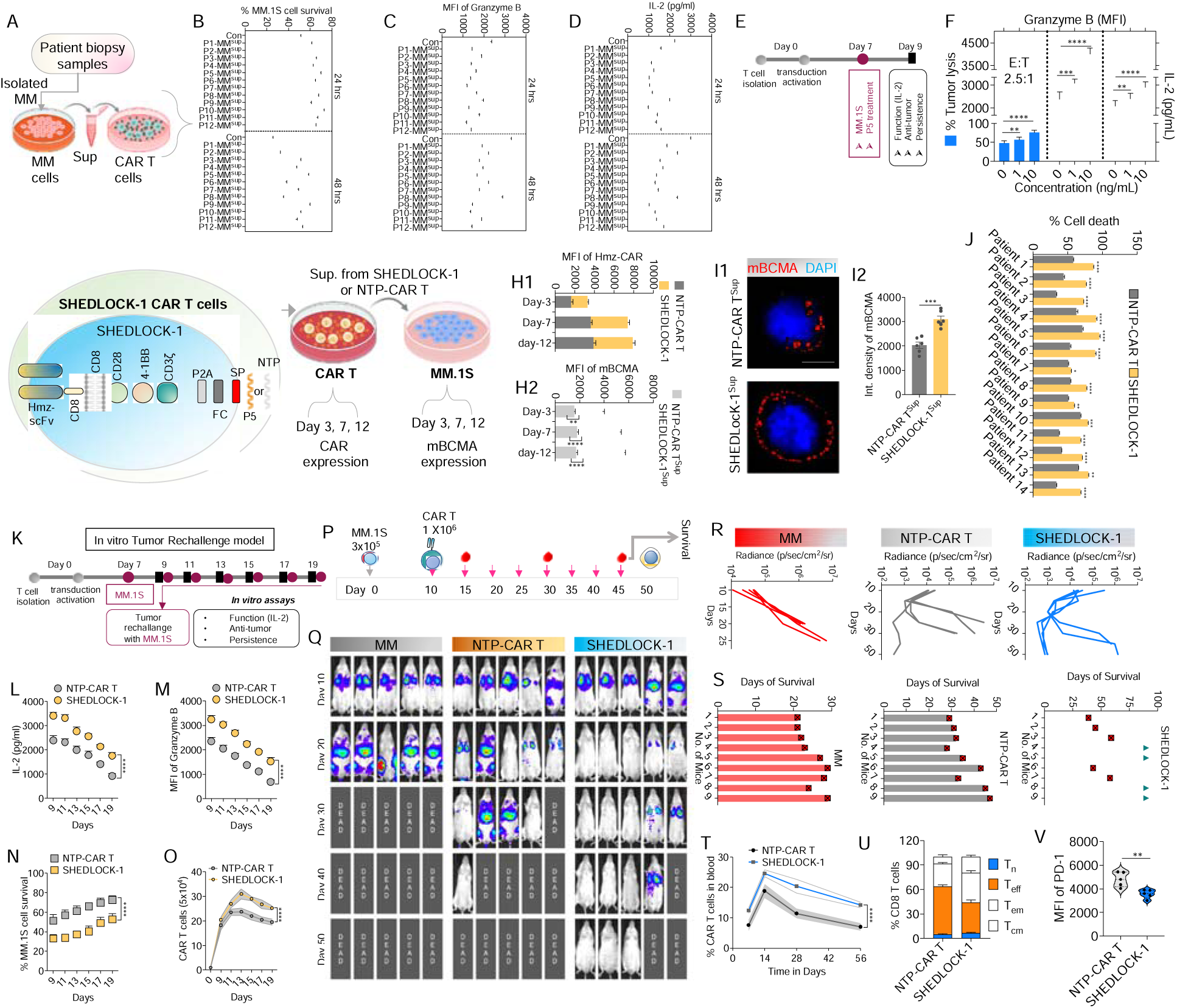
SHEDLOCK-1 CAR T cells boost anti-myeloma activity by blocking BCMA shedding. (A) Experimental setup showing supernatant collected from MM cells isolated from the patient biopsy samples (n=12 patient samples). CAR T cells were incubated with supernatant collected from MM cells and cultured in 1:1 ratio. (B) Bioluminescence-based cytotoxicity assay of MM.1S cells at two different time points upon co-culture with CAR T cells treated in the presence of MM supernatant containing sBCMA decoys. (C-D) Granzyme B and IL-2 assessment. Granzyme B was measured by flow cytometry and represented as MFI and IL-2 levels were measured by ELISA. (E) Illustration of the experimental setup showing CAR T cells treated with P5 peptide and co-cultured with MM.1S cells. (F) Bar graph showing tumor killing of MM.1S cells upon co-culture with CAR T cells and treated with different concentrations of P5. The bar graph also shows Granzyme B levels and IL-2 under similar conditions (n=5). Adding peptide P5 restores function in a dose-dependent manner and enhances killing. (G) Design of SHEDLOCK-1 CAR T cells expressing a secretory P5 or NTP (NTP-CAR T) molecule within CAR design. The CAR T cells were named as SHEDLOCK-1. Supernatant (sup) was collected from the SHEDLOCK-1 or NTP- CAR T cells at different time intervals after the transduction and this sup. was used to incubate the MM.1S cells. Non-targeting peptide (NTP) was used as control. (H1) MFI of the CAR T cells showing the transduction at various time points. (H2) Bar graph of flow cytometry analysis of MM.1S cells exposed to SHEDLOCK-1 sup shows increased mBCMA (reduced shedding). (I1) Super-resolution imaging validates enhanced mBCMA (red) retention. DAPI (blue) stained the nucleus. (I2) Bar graph of the image analysis (n=6). (J) Percentage of cell death in patient derived MM cells by NTP and SHEDLOCK-1 CAR T cells. Patient-derived MM cells are more efficiently cleared by SHEDLOCK-1 versus NTP-CAR (n=14 patient samples). (K) Illustration of the in vitro TR model. (L) Assessment of IL-2 levels at various time points upon TR (n=6). (M) Similarly, Granzyme B levels were measured by flow cytometry and presented as MFI (n=6). (N) Cytotoxicity as % of MM.1S cell survival measured by bioluminescence at different time points (n=6). (O) Number of CAR T cells analyzed over time upon TR, which shows persistence of these cells upon repeated tumor rechallenge. (P) MM.1S xenograft study schema and SHEDLOCK-1 or NTP-CAR T cell administration. (Q) Bioluminescence imaging (BLI) of the mice over time showing lower tumor burden and improved survival with SHEDLOCK-1 CAR T cell. (R) Image analysis showing kinetic pattern of the BLI signal over time (n=5). (S) Days of survival for individual mice, showing >60% survival at day 50 in SHEDLOCK-1 treated mice versus <20% in NTP-CAR T treated mice (n=9 mice per group). (T) CAR T cells measured in the blood over time (n=9). (U) Bar graph analysis of the flow cytometry data of the CAR T cell phenotype shows various T cells sub populations (n=5). (V) MFI of PD-1 represented as bar graph obtained from the flow cytometry data (n=5). \Data represent mean ± SEM. *p < 0.05; **p < 0.01; ***p < 0.001; ****p < 0.0001. A two-tailed Mann-Whitney U test was used for statistical analysis between groups, and a Two-way ANOVA followed by post-hoc testing was applied.

We then examined the cytolytic efficacy of SHEDLOCK-1 CAR T cells by measuring tumor lysis and cytokine production. Compared with NTP-CAR T cells, SHEDLOCK-1 CAR T cells showed clearly greater activity and cytotoxicity against MM .1S and MM.1R cells (**Supplementary Figure S8A -C**) and strong responses across multiple BCMA-positive targets (**Supplementary Figure S8D and S8E**). In a separate panel of 14 patient-derived MM samples, SHEDLOCK-1 CAR T cells killed tumor cells more effectively than NTP-CAR T cells (**Figure 3J**). In a long-term tumor-rechallenge (TR) assay, SHEDLOCK-1 CAR T cells sustained superior killing and secreted more Granzyme B and IL-2 (**Figure 3 K to 3M**). These advantages corresponded to better tumor control and longer persistence than controls (**Figure 3N and 3O**).

We evaluated in vivo efficacy in an MM.1S xenograft model (**Figure 3P**). Mice given SHEDLOCK-1 CAR T cells showed a sustained drop in tumor burden **(Figure 3Q and 3R)** and more than 60% survival at day 50, compared with less than 20% in control groups (n = 9 mice per group; **Figure 3S**). These effects came with durable CAR T cell persistence, an expanded central-memory pool, and lower expression of exhaustion markers **(Figure 3 T to 3 V).** To confirm that P5 secretion drove the benefit, we engineered non-BCMA-targeting CD19 CAR T cells to co-express P5 or NTP (**Supplementary Figure S9A**). In co-engrafted MM.1S and CD19-positive NALM6 models, only CD19-P5 CAR T cells lowered circulating and bone marrow sBCMA and promoted mBCMA retention on MM cells, whereas CD19-NTP controls did not (**Supplementary Figure S9B to S9D**). These results confirm that the strong anti-myeloma activity of SHEDLOCK-1 CAR T cells stems from effective inhibition of BCMA shedding.

Finally, we assessed safety in a cytokine release syndrome (CRS) model (**Supplementary Figure S10A**). SHEDLOCK-1 CAR T cells caused moderate increases in IL-1β, GM-CSF, TNF-α, IFN-γ, IL-2 and IL-6 (**Supplementary Figure S10B-G**), and liver function analyses showed modest, transient elevations in ALP, ALT, AST and albumin relative to NTP-CAR T controls, all of which remained below the levels measured in untreated MM.1S mice (**Supplementary Figure S10H-K**). Tocilizumab (TCZ) co-administration reduced the CRS-associated cytokine rises and restored organ function without impairing CAR T persistence or efficacy, as shown by maintained long-term survival and detectable CARnT cells (**Supplementary Figure S10-S11**). Overall, SHEDLOCK-1 CAR T cells show markedly greater cytotoxicity, persistence, and cytokine function than conventional CAR T cells. The accompanying CRS-like effects were mild and manageable and likely reflected secreted peptide activity or faster tumor lysis, and these findings support the translational promise of SHEDLOCK-1 as a next-generation, self-regulating CAR T platform for MM.

### A γ-secretase inhibiting peptide suppresses BCMA shedding and enhances CAR T cell activity

Beyond targeting the γ-secretase binding site on BCMA directly, we explored the natural substrates of γ-secretase, amyloid precursor protein (APP) and Notch, whose transmembrane regions are cleaved by the enzyme and therefore occupy its catalytic site^10, 12^. Our aim was to find endogenous or semi-synthetic sequences that lower immunogenicity and can serve as a secreted regulatory module within the CAR constructs. Following this rationale, we selected peptides derived from the well-characterized γ-secretase substrates APP and Notch (**Figure 4A**). We modeled Notch, APP, and their analogues (nx-P1 and nx-P2) with the modified residues shown in cartoon representation **(Figure 4B** **and Supplementary Figure S12)**. We also modeled the γ-secretase complex, made up of Nicastrin (NCT), Presenilin-1 (PS-1), Presenilin enhancer-2 (PEN-2), and APH-1, bound to these substrates, and we designed the site-directed variants nx-P1 and nx-P2 by altering the γ-secretase cleavage motifs. This modeling identified PS-1 as the main catalytic subunit that recognizes and cleaves substrate **(Figure 4E1).** MD simulations of PS-1 bound to Notch, APP, nx-P1, and nx-P2 within POPC: DOPC bilayers showed distinct stability throughout the trajectories, and MM/GBSA analysis showed that the binding energy profiles were similar or better for the peptide substrates, especially the modified Notch-derived analogues **(Figure 4C1, 4C2 and 4D; Movie 2; Supplementary Figure S13 and** **Table 2).** Representative snapshots captured time-dependent conformational changes in PS-1 and the bound peptide, consistent with altered substrate engagement and complex stability **(Figure 4E2).** All peptide-PS-1 complexes maintained stable interactions within the catalytic pocket, supporting their potential to act as competitive γ-secretase inhibitors.

**Figure 4:**
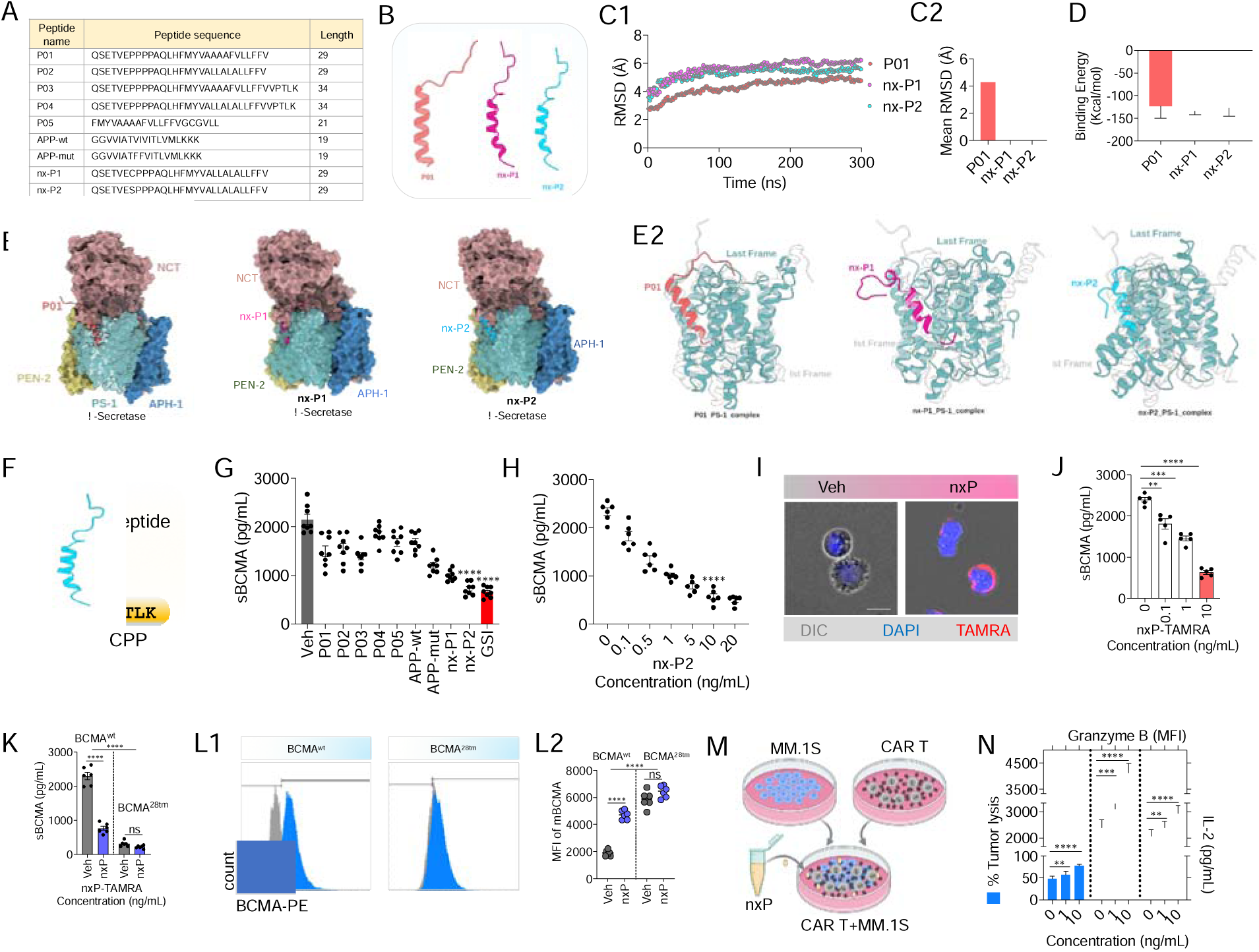
Endogenous γ-secretase-interacting peptides suppress shedding and enhance CAR function. (A) Table showing the list of the peptides that were screened. (B) Structural models of P01, nx-P1 and nx-P2. (C1, C2) RMSD trajectories and mean RMSD for the P01, nx-P1 and nx-P2 complexes with PS-1. (D) MM/GBSA binding free energies for the same complexes. (E1) Surface representation of Presenilin-1-substrate complex models embedded within a POPC:DOPC bilayer (lipid bilayer not shown for clarity), illustrating the positioning of each peptide within the γ-secretase catalytic groove. (E2) Superposition of the first and last simulation frames for each peptide-PS-1 complex. MD simulations show that engineered Notch-derived peptide analogues maintain the most persistent interactions with the active-site pocket in PS-1 as depicted by their converged RMSD plots, with reduced lateral displacement and greater TM-helix engagement over time. Superposition of the snapshots from MD simulations did not reveal much structural deviation in the complex between Notch-analogues and PS-1. (F) Peptide synthesis workflow incorporating a cell-penetrating peptide (CPP) for efficient cell uptake. (G, H) ELISA assay of the sBCMA levels upon treatment with various peptides and evaluation of different concentrations of the shortlisted nx-P2 (designated nxP) peptide demonstrates the strongest suppression of γ-secretase–mediated BCMA cleavage. (I) Representative images of nxP-TAMRA (red) shown as signal accumulation at the plasma membrane. DIC image is shown in the grey (n=8 images). (J) Bar graph showing ELISA assay results of sBCMA in cells treated with nxP-TAMRA at various concentrations (n=5). (K) ELISA assay showing cleavage-resistant BCMA variant (BCMA-28^tm^, containing the CD28 transmembrane domain). nxP produces minimal reduction in sBCMA in BCMA-28^tm^ cells, confirming that its activity depends on the native BCMA γ-secretase cleavage motif. (L1-L2) Representative flow cytometry histograms of cells expressing wild type (wt) or BCMA-28^tm^ and the corresponding bar graphs (n=6). (M) Work flow of co-culture assays with BCMA CAR T cells and MM.1S targets upon treatment with nxP. (N) Bar graph of % tumor lysis, Granzyme B, and IL-2, indicating improved functional activation and killing efficiency upon treatment with increasing nxP peptide concentration (n=6). Data represent mean ± SEM. *p < 0.05; **p < 0.01; ***p < 0.001; ****p < 0.0001. A two-tailed Mann–Whitney U test was used for statistical analysis between groups.

We then synthesized these peptides with a membrane-penetrating CPP sequence to aid cellular uptake (**Figure 4F**) and screened them in MM.1S cells for inhibition of BCMA shedding. All peptides reduced sBCMA release, with significant inhibition by the engineered Notch-derived analogues nx-P1 and nx-P2 **(Figure 4G),** and a modified Notch-derived peptide (nx-P2) gave the strongest effect and was chosen for further study, hereafter called nxP (**Figure 4H**). To confirm intracellular localization, we labeled nxP with tetramethylrhodamine (TAMRA); nxP-TAMRA fluorescence localized mainly to the plasma membrane while retaining potent blockade of BCMA shedding (**Figure 4I and 4J**). To test whether nxP acted specifically through γ-secretase inhibition, we generated a cleavage-resistant BCMA mutant carrying the CD28 transmembrane domain (BCMA28tm). Consistent with a γ-secretase-dependent mechanism, nxP had little effect on mBCMA or sBCMA in BCMA28tm-expressing HEK293 cells but raised mBCMA and lowered sBCMA in wild-type BCMA cells **(Figure 4 K, 4L1 and 4L2)**. Mass spectrometry confirmed nxP in the cell supernatant of SHEDLOCK-2 cells (**Supplementary Figure S14**).

To see how γ-secretase inhibition affects CAR T activity, we tested nxP in co-cultures of CAR T cells with MM.1S targets. nxP increased CAR T cytotoxicity and raised secretion of Granzyme B and IL-2 relative to untreated controls (**Figure 4M and 4N**). nxP did not affect the proliferation or viability of several cell types, indicating specificity and no off-target toxicity (**Supplementary Figure S15**). Together, these findings show that nxP suppresses BCMA shedding through selective γ-secretase inhibition and strengthens the functional potency of CAR. T cells.

### CAR-integrated nxP secretion enhances CAR T potency and persistence

To further exploit peptide-mediated inhibition of BCMA shedding, we engineered CAR T cells to secrete the naturally derived nxP peptide as part of the CAR molecule using a single-plasmid construct that was then transduced into T cells (**Figure 5A**). We named these SHEDLOCK-2 CAR T cells and compared them directly with SHEDLOCK-1 CAR T cells, which secrete the synthetic P5 peptide. No significant differences were observed in transduction efficiency between SHEDLOCK-2 and SHEDLOCK-1; however, a slight reduction was noted compared to the hmz-CAR backbone, likely attributable to the lower vector titre resulting from the larger transgene size (**Supplementary Figure S15C-F**). To verify that the integrated nxP was functional, we first confirmed its secretion and inhibitory activity by demonstrating that supernatants collected from SHEDLOCK-2 CAR T cells at several time points after transduction stabilized mBCMA on MM cells **(Figure 5B1 and 5B2).** Having established nxP functionality, we next assessed cytotoxicity and found that SHEDLOCK-2 CAR T cells were more cytotoxic than SHEDLOCK-1 CAR T cells against a range of BCMA-expressing targets, demonstrating that nxP secretion not only preserved but enhanced tumor killing **(Figure 5C** **and Supplementary Figure S16C to S16).** Treating BCMA-positive tumor lines with nxP (20 ng/mL) did not affect apoptosis **(Supplementary Figure S16F)**, ruling out direct tumor-intrinsic toxicity. Continuous nxP exposure slightly reduced CAR T proliferation (**Figure 5D**), which prompted us to test long-term function in an in vitro TR model (**Supplementary Figure S17A**). SHEDLOCK-2 CAR T cells showed stronger anti-tumor activity and better in vitro persistence, consistent with enhanced proliferation on repeated antigen exposure **(Supplementary Figure S17B and S17C).** Immunophenotyping revealed a higher proportion of central memory T cells in SHEDLOCK-2 CAR T cells, revealing their greater durability and indicating that CAR-integrated nxP secretion favors a less-differentiated memory phenotype that is able to sustain repeated antigen encounters **(Supplementary Figure S17D1 and S17D2).**

**Figure 5:**
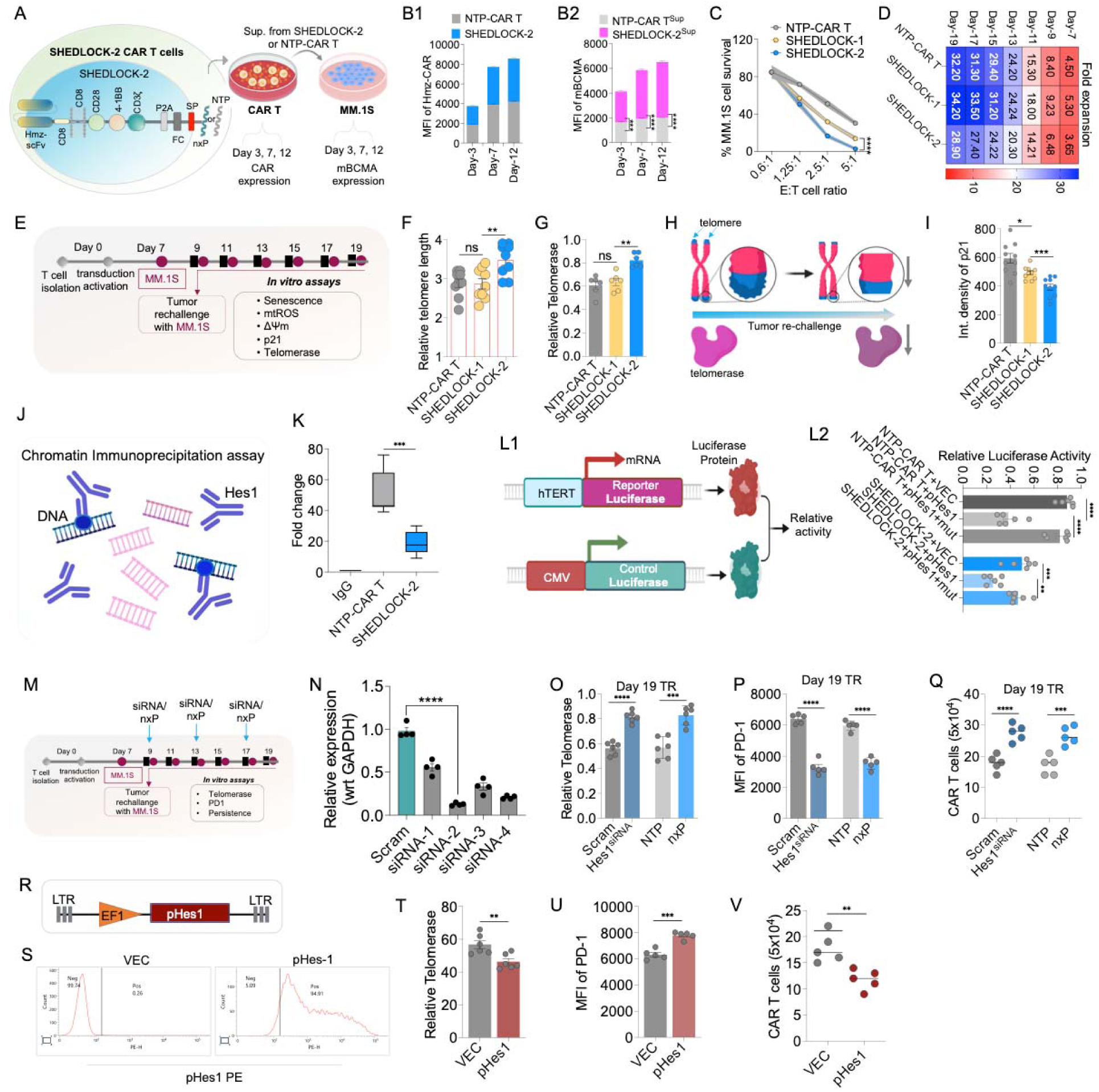
Telomere preservation and reduced senescence underpin the persistence of SHEDLOCK-2 CAR T cells. (A) Schematic showing the design of SHEDLOCK-2 CAR T cells with secretory Notch-modulatory peptide nxP. The supernatant obtained from SHEDLOCK-2 CAR T cells at various time points after CAR T transduction is used to incubate MM.1S cells with 1:1 ratio of cell culture media. (B1) CAR T transduction efficiency of SHEDLOCK-2 CAR T cells and NTP-CAR T cells over time and (B2) MFI of mBCMA on MM.1S cells cultured in presence of the sup. obtained from SHEDLOCK-2 CAR T cells and NTP-CAR T Cells (n=6). (C) The cytotoxic assay done by bioluminescence showing that SHEDLOCK-2 CAR T cells has more robust anti-tumor activity than SHEDLOCK-1 CAR T cells (n=5). (D) Proliferation rate shown as heat map of the cells and represented as fold expansion. The assay was performed by automated cell counter (n=10). (E) Workflow of in vitro TR assay and the assay parameters at the end of the experiment. (F) Relative telomere length of SHEDLOCK-1 and SHEDLOCK-2 CAR T cells with NTP-CAR T cells used as controls upon TR analysed by qPCR where SHEDLOCK-2 CAR T cells showed enhanced telomere length compared with SHEDLOCK-1 and NTP-CAR T cells (n=9). (G) Relative telomerase activity analysis by qPCR in SHEDLOCK-2 vs SHEDLOCK-1 CAR T Cells with NTP-CAR T cells used as controls (n=6). (H) Schematic of the telomere length assay showing telomere shortening in cells upon tumor rechallenge. (I) Image analysis of the cells stained with anti-p21 antibody to determine the p21 signal. The images were captured by the microscopy and quantified using ImageJ and represented as integrated (int.) density (n=10). (J) Schematic illustration of the chromatin immunoprecipitation (ChIP) assay used to evaluate sequential TR for assessment of telomerase activity, PD-1 expression, and CAR T cell persistence. (K) ChIP-qPCR analysis showing Hes1 enrichment at the hTERT promoter in NTP-CAR T vs SHEDLOCK-2 CAR T Cells. (L1) Schematic of the dual-luciferase reporter assay used to quantify hTERT promoter activity following modulation of Hes1 signaling. (L2) Relative hTERT promoter activity measured by dual-luciferase reporter assay following Hes1 overexpression or mutant Hes1 expression in NTP-CAR T and SHEDLOCK-2 CAR T cell (n=6). (M) Experimental workflow illustrating siRNA-mediated Hes1 knockdown during (N) Validation of Hes1 siRNA knockdown efficiency by quantitative RT-PCR relative to GAPDH (n=4).Hes1 occupancy at the hTERT promoter. (O) Relative telomerase activity in CAR T cells following Hes1 knockdown or nxP treatment at Day 19 tumor rechallenge. (P) Flow cytometric quantification of PD-1 expression following Hes1 knockdown or nxP treatment at Day 19 tumor rechallenge. (Q) Bar graph showing persisting CAR T cells after Hes1 knockdown or nxP treatment at Day 19 tumor rechallenge. (R) Schematic representation of the lentiviral EF1α-pHes1 expression construct used for Hes1 overexpression studies. (S) Representative flow cytometry histograms confirming Hes1 overexpression following lentiviral transduction. (T) Relative telomerase activity following ectopic Hes1 overexpression in CAR T cells. (U) Flow cytometric analysis of PD-1 expression following Hes1 overexpression. (V) Quantification of CAR T cell persistence following Hes1 overexpression. Data represent mean ± SEM. *p < 0.05; **p < 0.01; ***p < 0.001; ****p < 0.0001. A two-tailed Mann–Whitney U test was used for statistical analysis between groups.

### Telomere preservation and reduced senescence enhance persistence of SHEDLOCK-2 CAR T cells

To understand why nxP improved CAR T persistence, we first profiled cell metabolism and found that SHEDLOCK-2 CAR T cells favored oxidative phosphorylation (OXPHOS) over glycolysis, a hallmark of long-lived memory T cells^25, 43^ **(Supplementary Figure S17E1, S17E2, S17F1 and S17F2)**. In agreement, flow cytometry showed a decrease in the exhaustion markers PD-1, TIM-3, and LAG-3 compared with SHEDLOCK-1 CAR T cells (**Supplementary Figure S18**). For a broader comparison, we benchmarked SHEDLOCK-2 CAR T against a second-generation BCMA CAR T construct of the type used in ide-cel. Comparatively, SHEDLOCK-2 cells showed robust activity against patient-derived MM cells with greater CAR T persistence, adding clinical context for its therapeutic advantage (**Supplementary Figure S19**). RNA-seq comparison of NTP-CAR T cells and SHEDLOCK-2 CAR T cells revealed enrichment of metabolic pathways including glycolysis, OXPHOS, and FOXO signaling (**Supplementary Figure S20**), pointing to broad metabolic reprogramming behind SHEDLOCK-2 CAR T persistence and function.

Given the known role of Notch signaling in cellular senescence and T cell exhaustion, we next examined telomere dynamics and senescence markers in SHEDLOCK-2 CAR T cells, especially as recent work shows that CAR T cells undergo telomere shortening and mitochondrial dysfunction after infusion and tumor re-challenge^24, 44^. In the in vitro TR model, nxP, but not P5, reduced senescence and restored mitochondrial (mt) ROS levels and membrane potential (**Supplementary Figure S21**). Measuring telomerase activity and telomere length in the TR model, we found that SHEDLOCK-2 CAR T cells preserved both, whereas these declined in NTP-CAR T cells (**Figure 5E to 5H**). In line with the connection between telomere dysfunction and p21 activation, SHEDLOCK-2 cells showed markedly lower p21 expression (**Figure 5I**). To test whether telomere maintenance by nxP works through Notch signaling, we used our reporter cell-line system and found that nxP inhibited Notch cleavage in a dose-dependent manner, which was associated with higher telomerase activity and telomere retention **(Supplementary Figure S22A to S22C).** Incubating CAR T cells with nxP also reduced NICD generation, indicating that nxP specifically blocks γ-secretase-mediated cleavage of Notch (**Supplementary Figure S22D**).

### Hes1 represses hTERT transcription downstream of Notch to limit telomerase activity and CAR T persistence

To find how Notch inhibition lowers telomerase activity, we examined the hTERT promoter and identified three strong Hes1 binding sites within the 2 kb region upstream of the transcription start site, located near -579 (GGCACGCG, contains CACGCG class C sites having strong Hes1 compatible site), -282 (GGCGCGCG, CG-rich Hes1 prediction), and -163 bp (GCCACGTG, canonical E box, CACGTG). (**Supplementary Figure S23**). Chromatin immunoprecipitation (ChIP)-qPCR with an anti-Hes1 antibody confirmed this enrichment in NTP-CAR T cells, which fell substantially in SHEDLOCK-2 cells (**Figure 5J and 5 K**). Using an hTERT proximal promoter-luciferase reporter, we found that co-expression of Hes1 repressed promoter activity by about 50%, and this repression disappeared when the three predicted Hes1 sites were mutated, showing that these sites are required for Hes1-mediated repression **(Figure 5L1 and 5L2).**

We also measured Hes1 expression under in vitro TR conditions. Over time, Hes1 rose significantly in NTP-CAR T cells but remained significantly lower in SHEDLOCK-2 CAR T cells, and this reduced expression coincided with restored telomerase activity relative to NTP CAR T cells (**Supplementary Figure S24**). The rise in Hes1 was accompanied by greater exhaustion and reduced persistence, both of which were restored under SHEDLOCK-2 conditions (**Supplementary Figure S24**). Under TR conditions, lowered exhaustion, and increased persistence, reproducing the nxP effect seen in SHEDLOCK-2 CAR T cells (Hes1 siRNA rescued telomerase activity, **Figure 5M to 5Q**). Conversely, Hes1 overexpression in resting CAR T cells was sufficient to suppress hTERT and telomerase, raise exhaustion markers, and sharply reduce CAR T cell persistence (**Figure 5R to 5 V)**.

To confirm that Hes1 expression depends on Notch activation, we used an shRNA to knock down Notch1. Notch1 knockdown markedly lowered Hes1 expression after TR and increased telomerase activity (**Supplementary Figure S25**). Overexpressing Hes1 in Notch1-knockdown CAR T cells reduced telomerase activity and induced exhaustion with fewer durable CAR T cells (**Supplementary Figure S25**). To determine whether the benefit of nxP comes specifically from blocking Notch signaling rather than from broader γ-secretase inhibition, we generated a lentiviral construct encoding NICD1-ΔE, a constitutively nuclear, γ-secretase-independent form of the Notch1 intracellular domain, and transduced it into SHEDLOCK-2 CAR T cells (**Supplementary Figure S26A**). Despite ongoing nxP secretion, NICD1-ΔE restored Hes1 expression, and telomerase activity, and returned p21 to NTP-CAR T levels (**Supplementary Figure S26B-D**). Importantly, when tested against BCMA28OO tumor cells-both NTP and SHEDLOCK-2 cells showed similar anti-tumor activity and no further protective effect on preserving telomerase activity/length, suggesting that the effect of nxP on telomere maintenance is specifically mediated by Notch signaling inhibition (**Supplementary Figure S26E-H**). This downstream reconstitution shows that the benefit of nxP mainly comes from inhibiting Notch transcriptional output rather than from effects on other γ-secretase substrates.

### SHEDLOCK-2 CAR T cells show robust anti-tumor activity and long-term persistence in vivo

To test whether the naturally derived γ-secretase inhibitory peptide nxP could improve CAR T function in vivo, we established an MM xenograft model in NSG mice (**Supplementary Figure S27A**). We gave SHEDLOCK CAR T cells on day 10 after MM engraftment and recovered CAR T cells from treated mice on day 14. We then cultured the recovered cells ex vivo and applied supernatants from SHEDLOCK-1 and SHEDLOCK-2 CAR T cells to MM.1S cells to measure inhibition of sBCMA release. The sBCMA analysis confirmed effective expression and secretion of the γ-secretase-interacting peptides P5 and nxP, and supernatant from nxP-expressing SHEDLOCK-2 CAR T cells suppressed sBCMA far more strongly than SHEDLOCK-1 supernatant. These data validate successful in vivo engineering, secretion competency, and functional activity of the peptide-secreting CAR T platforms (**Supplementary Figure S27B**).

We next developed a further xenograft model to evaluate SHEDLOCK CAR T cells in depth (**Figure 6A**). At two doses, SHEDLOCK-2 CAR T cells produced potent anti-tumor activity (**Figure 6B**), and even the lowest dose cleared tumor as well as the higher dose and gave durable disease control. Longitudinal monitoring showed that tumor regression lasted more than 120 days (last data point; group 1, n = 10 mice) and showed a clear improvement in survival (**Figure 6C**). In vivo tracking detected SHEDLOCK-2 CAR T cells in both blood and bone marrow (**Figure 6D and 6E)** confirming sustained engraftment and long-term durability (group 2, sacrificed at day 50). Both P5 and nxP appeared in blood at early time points and declined over time, consistent with the decline in CAR T cells, and both were detected in bone marrow at levels that matched the CAR T cells in these compartments. Quantification with custom sandwich ELISAs showed that P5 and nxP concentrations in peripheral blood peaked at day 15 (LD: ∼10–20 pg/mL; HD: ∼44-53 pg/mL), declined to ∼2-15 pg/mL by day 30, and fell below the limit of detection (≤0.5 pg/mL) by day 45, paralleling CAR T cell contraction. In contrast, bone marrow aspirates at day 50 retained substantially higher concentrations (LD: ∼5-12 pg/mL; HD: ∼20-55 pg/mL), confirming preferential peptide accumulation within the tumor microenvironment (**Supplementary Figure S28**). Together, these results show that SHEDLOCK-2 CAR T cells stay effective even at reduced doses, supporting their therapeutic robustness.

**Figure 6:**
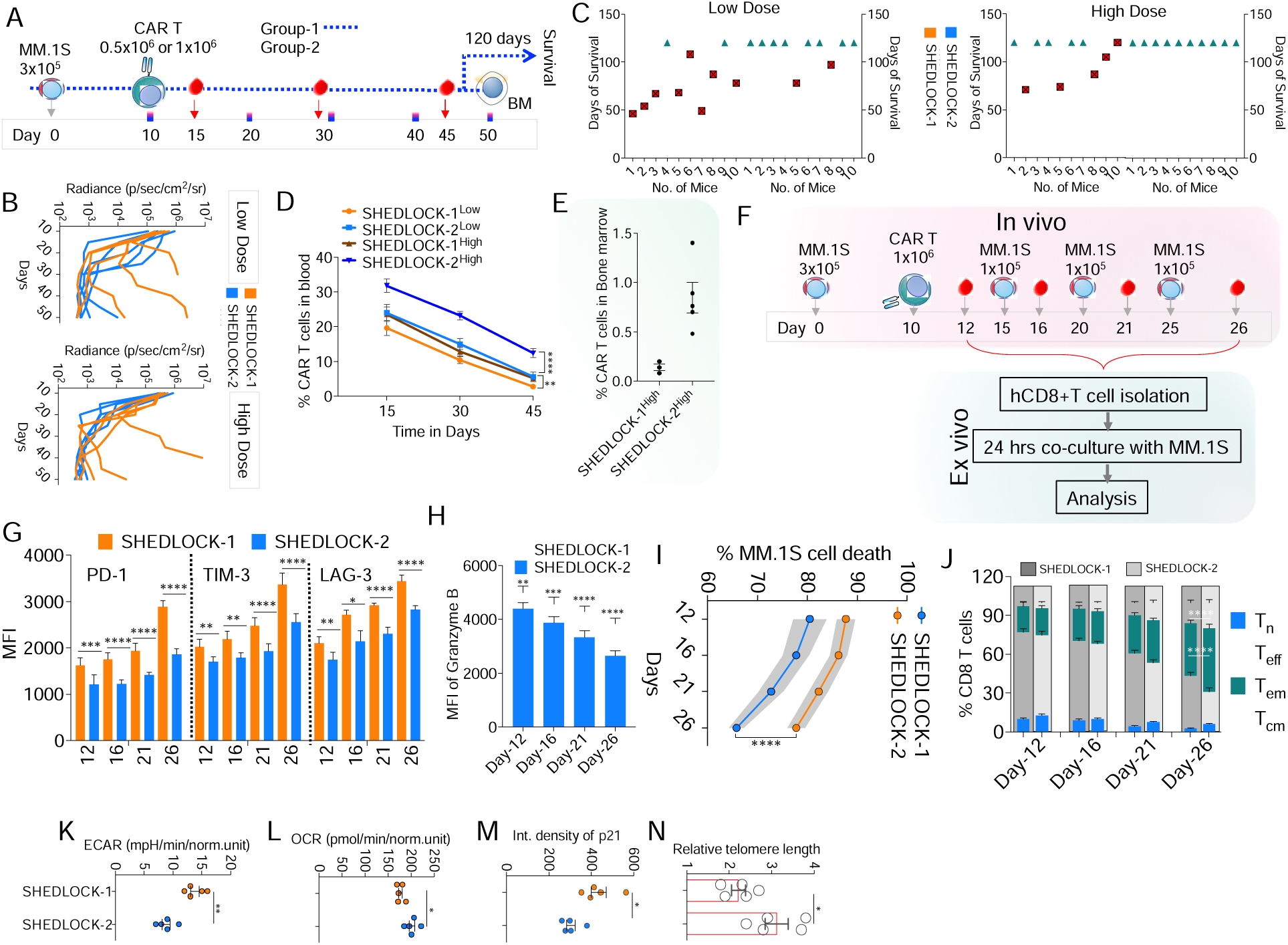
SHEDLOCK-2 CAR T cells sustain potent in-vivo activity and long-term persistence. (A) Schematic of the MM.1S xenograft study design showing tumor cell injection followed by the administration of SHEDLOCK-2 CAR T cells at two dose levels (0.5×10^6^ or 1×10^6^ cells). Time points for imaging, blood collection, and endpoint analyses are indicated in the schema. The mice were divided into two groups; group-1 was continued for survival kinetics and group-2 mice were sacrificed at day 50 for end point analysis. (B) BLI of MM.1S tumor-bearing mice treated with low-dose or high-dose SHEDLOCK-2 CAR T cells across the indicated study days showing quantification of the BLI signal as radiance (p/sec/cm²/sr) measured at the corresponding time points (n=5). (C) Days of survival for individual mice in the low-dose and high-dose groups (n=10 mice per group). (D) Graph of flow cytometry quantification of circulating CAR T cells isolated from peripheral blood at sequential time points following infusion (n=5). (E) Percentage of CAR T cells detected in bone marrow at day 50, quantified by flow cytometry (n=5). (F) Schematic of the experimental workflow of CAR T cells harvested in the TR mice at various time intervals. These CAR T cells were then evaluated ex vivo. (G) Bar graph showing the MFI of exhaustion markers, PD-1, TIM-3 and LAG-3 at various time points (n=5). (H) Representative plots and bar-graph quantification of Granzyme B measured by flow cytometry and represented as MFI for SHEDLOCK-1 and SHEDLOCK-2 CAR T cells (n=5). (I) Anti-tumor activity as measured by the bioluminescence at different time points (n=5). (J) Immunophenotyping of the various CAR T cell subsets showing naïve T cells (T_n_); effector T cells (T_eff_), central memory (T_cm_), and effector memory (T_em_) cell population (n=5). (K, L) Bar graph representing expression of mitochondrial activity showing ECAR and OCR (n=5). (M) Image analysis of p21 immunofluorescence was performed using ImageJ software, and signal intensity was quantified as integrated density. (n=5). (N) Relative telomere length quantification done by qPCR (n=5). Data represent mean ± SEM. *p < 0.05; **p < 0.01; ***p < 0.001; ****p < 0.0001. A two-tailed Mann–Whitney U test was used for statistical analysis between groups. For the longitudinal data in panels G to J, a two-way ANOVA followed by post-hoc testing was applied.

To assess functional durability under tumor re-challenge, we established a TR model, isolated single cells from TR mice at defined intervals, and analyzed CAR T phenotype and function (**Figure 6F**). Flow cytometry showed lower exhaustion markers (PD-1, TIM-3, and LAG-3) in SHEDLOCK-2 than in SHEDLOCK-1 CAR T cells (**Figure 6G**) along with sustained Granzyme B secretion (**Figure 6H**). SHEDLOCK-2 CAR T cells kept their anti-tumor activity over time and contained a higher fraction of memory-phenotype cells than SHEDLOCK-1 cells (**Figure 6I and 6J**). Metabolic profiling reinforced this pattern, showing a shift toward OXPHOS over glycolysis that reflects a long-lived memory state (**Figure 6 K and 6L**). SHEDLOCK-2 CAR T cells also expressed far lower senescence-associated p21 **(Figure 6M)** and maintained telomere length **(Figure 6N).** Together, these findings show that the nxP-based SHEDLOCK-2 construct drives coordinated metabolic and transcriptional reprogramming that supports memory formation, mitochondrial fitness, and telomere preservation, and thereby extends CAR T cell longevity.

### CAR T cell metabolic activity and telomere function predict efficacy outcome

Because telomere function, measured as telomerase activity and telomere length, declines over time under tumor re-challenge and is maintained by Notch inhibition, we developed an in vivo tumor re-challenge model using T cells from patients and measured mitochondrial OCR and ECAR, telomere length, Hes1 expression, CAR T persistence, and anti-tumor activity. We assessed these parameters before infusion and over time after re-challenge. TR reduced the mitochondrial metabolic parameters together with telomerase activity and CAR T persistence, while Hes1 expression and T cell exhaustion both rose **(Supplementary Figure S29).** To test whether these parameters predict clinical outcome, we extended the analysis to CAR T cells from 15 patients and generated SHEDLOCK-2 cells (**Supplementary Figure S30**). Telomere length and telomerase activity correlated strongly and positively with CAR T persistence and mouse survival and negatively with exhaustion (**Supplementary Figure S30**). Hes1 expression correlated negatively with persistence and survival and positively with exhaustion (**Supplementary Figure S30**). Mitochondrial metabolic parameters also correlated well with persistence, exhaustion, and survival. Overall, these preliminary mouse studies suggest that telomere length, Hes1 expression, and mitochondrial metabolic parameters correlate strongly with long-term CAR T function and efficacy in this preclinical model, and warrant prospective validation as candidate biomarkers.

### SHEDLOCK-2 CAR T cells show potent in vivo activity in serial transfer and patient-derived MM models

Because SHEDLOCK-2 CAR T cells retained stem-like features and showed reduced senescence in vitro, we next tested in vivo efficacy in a serial transfer model (**Figure 7A**). We isolated CAR T cells from primary NSG recipients and transferred them into secondary MM-bearing NSG mice. Ten days after infusion, we harvested cells from spleen and bone marrow and enriched CD3-positive cells with magnetic beads, confirming sustained CAR expression (hereafter referred to as serially transferred CAR T cells, or CAR T-ST; Figure 7B1 and 7B2). Each secondary recipient received 0.3 × 10^6^ MM cells and 0.5 × 10^6^ CAR T-ST cells. This adoptive transfer produced robust tumor suppression and significantly improved survival (**Figure 7C to 7E**). Among the groups, SHEDLOCK-2 CAR T-ST cells showed the strongest anti-tumor activity (**Figure 7C and 7D**) and the longest survival (n = 10 mice per group) relative to NTP and SHEDLOCK-1 CAR T-ST cells (**Figure 7E**). SHEDLOCK-2 CAR T-ST cells stayed detectable throughout the study (**Figure 7F**), indicating durable persistence and continued tumor surveillance. These results confirm that SHEDLOCK-2 CAR T cells retain functional potency after in vivo expansion and show enhanced anti-myeloma activity in a clinically relevant system. To further validate efficacy, we developed a patient-derived multiple myeloma (PDx-MM) model based on a previously described protocol^45^, with modifications, using busulfan- conditioned NSG mice supplemented with human IL-6 (NSG-exIL6; **Figure 7G**). These mice showed characteristic features of myeloma progression, with human Ig detected in plasma at regular intervals by ELISA (**Supplementary Figure S31**) and progressive human Ig accumulation in urine (**Supplementary Figure S32**). We then tested SHEDLOCK-2 CAR T cells in this model. Two weeks after infusion, serum human Ig fell substantially in all CAR T-treated groups, accompanied by detectable circulating CAR T cells (**Figure 7H**). Over time, SHEDLOCK-2-treated mice showed near-complete tumor clearance, undetectable Ig, fewer MM cells in blood, and improved survival (**Figure 7I**; **Supplementary Figures S31, S32, and S33**). SHEDLOCK-2 CAR T cells persisted in blood for up to six weeks after treatment (**Figure 7I**), confirming long-term durability and efficacy against patient-derived MM cells. SHEDLOCK-2 treatment also significantly reduced bone marrow kappa light chain (Igκ-positive) MM cells across all patient-derived samples, as shown by flow cytometry (**Supplementary Figure S34**).

**Figure 7:**
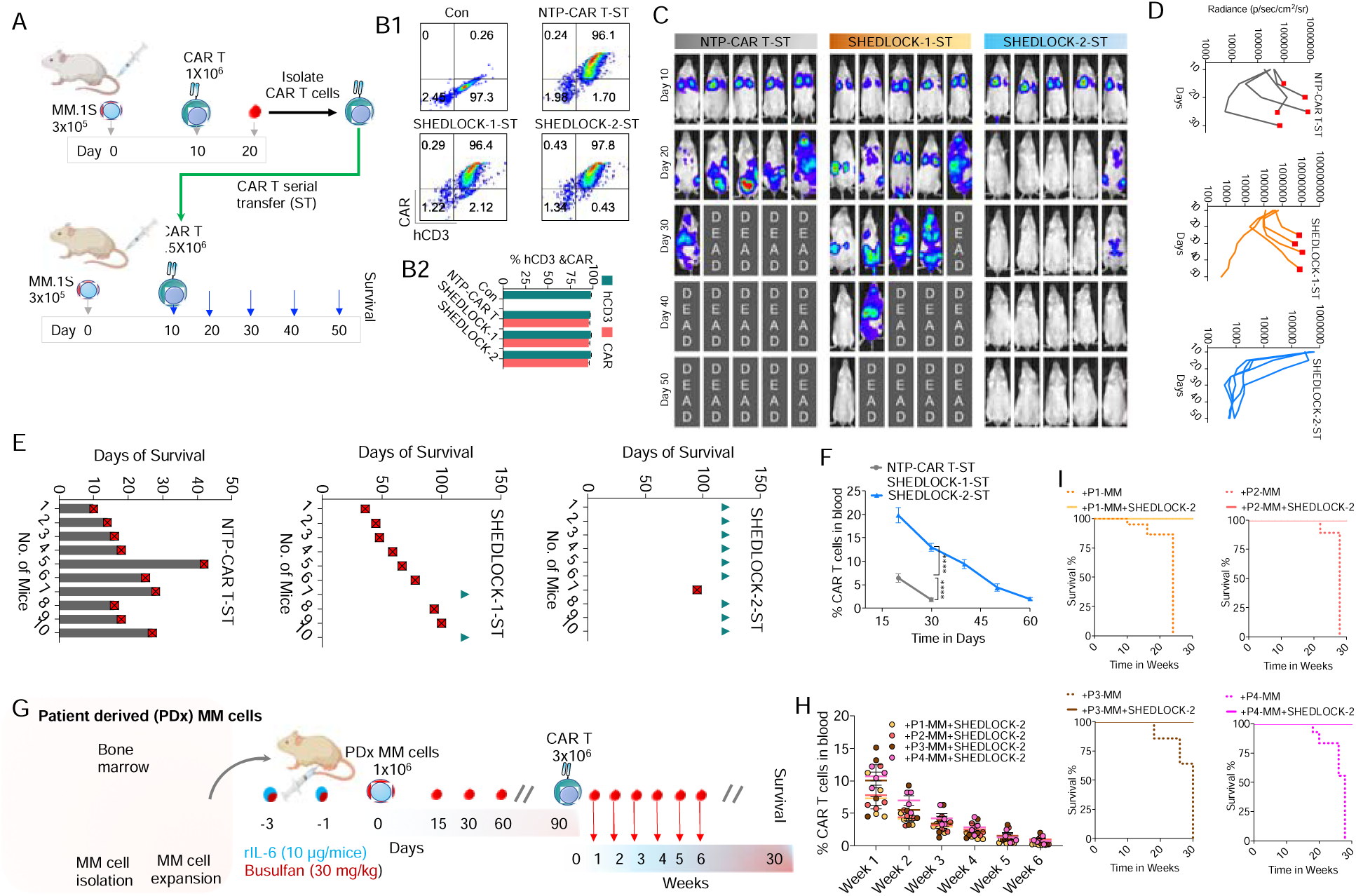
SHEDLOCK-2 remains effective after serial transfer and in a patient-derived myeloma model. (A) Workflow schematic outlining the experimental setup used to evaluate SHEDLOCK-2 CAR T cell phenotype, functional activity, and imaging-based readouts under serial transfer experimental setup. (B1) Representative flow cytometry plots showing hCD3 and CAR expression in serially transferred NTP-CAR T, SHEDLOCK-1 and SHEDLOCK-2 CAR T cells, with untransduced cells as control. (B2) Corresponding bar graph quantifying the percentage of hCD3-positive and CAR-positive cells (n=5). (C) BLI images of the mice over time showing lower tumor burden and improved survival with SHEDLOCK-2 CAR T cells. (D) Image analysis showing kinetic pattern of the BLI signal over time (n=5). (E) Days of survival for individual mice, showing the longest survival in SHEDLOCK-2 CAR T-ST treated mice compared with SHEDLOCK-1 or NTP-CAR T-ST treated mice (n=10 mice per group). (F) CAR T cells measured in the blood over time (n=5). (G) Workflow of the patient derived (PDx) MM cells and injected into the NSG mice (n=4 mice per group). The NSG mice also received Busulfan (30 mg/kg) and recombinant IL6 (rIL6). Various parameters were measured as indicated. (H) Flow cytometry analysis of the CAR T cells obtained from the blood at indicated time points. (I) Survival kinetics of the mice received MM cells from different patients (P1, P2, P3 and P4) followed by SHEDLOCK-2 treatment. Data represent mean ± SEM. *p < 0.05; **p < 0.01; ***p < 0.001; ****p < 0.0001. A two-tailed Mann–Whitney U test was used for statistical analysis between groups. For the longitudinal data in panels F and H, a two-way ANOVA followed by post-hoc testing was applied; survival in panels E and I was compared using the log-rank (Mantel-Cox) test.

### SHEDLOCK-2 CAR T cells show a favorable safety profile

To evaluate safety, we first tested SHEDLOCK-2 CAR T cells in a CRS model (**Supplementary Figure S35A**). Serum cytokine profiling showed no disproportionate rise in IL-6, IFN-γ, GM-CSF, IL-1β, TNF-α or IL-2; each was significantly lower in SHEDLOCK-2-treated than in SHEDLOCK-1-treated mice (**Supplementary Figure S35B to S35F**). Peritoneal immune monitoring showed no expansion of monocyte, macrophage, neutrophil, eosinophil or dendritic cell populations beyond what tumor regression would explain **(Supplementary Figure S35G to S35 K).** Biochemical analyses confirmed that hepatic and renal parameters stayed within physiological ranges, with no sign of systemic organ toxicity (**Supplementary Figure S35L to S35R**). Histopathology supported these findings, showing no necrosis or inflammation in major organs apart from minor background changes linked to tumor clearance (**Supplementary Figure S36**).

To extend the safety assessment to later time points, we next developed a sub-chronic model over 90 days using NCG mice and tested SHEDLOCK-2 cells alongside a sham control. Because many SHEDLOCK-1 mice relapse by day 90, we used only SHEDLOCK-2 to evaluate extended safety. Assessment of IL-10, IL-17A, IL-12p70, and MCP-1, in addition to the original IL-6, TNF-α, IL-1β, IL-2, GM-CSF, and IFN-γ, revealed no significant changes, and the values were close to those of sham control mice (**Supplementary Figure S37**). We also performed complete blood counts to assess hematopoiesis, cardiac troponin-I as a sensitive marker of cardiac injury, cystatin-C together with creatinine and BUN for renal function, and ALT, AST, ALP, and albumin for hepatic function (**Table 3**). A blinded, board-certified pathologist re-evaluated histopathology of liver, kidney, lung, heart, spleen, intestine, brain, and bone marrow with semi-quantitative scoring on a 0 to 4 scale and identified no drug-related lesions in any organ system (**Supplementary Figure S38 &** **Table 4**).

Together, these results show that SHEDLOCK-2 CAR T cells produce potent anti-myeloma activity while maintaining an acceptable safety profile, with inflammatory responses staying within manageable limits. These findings highlight the translational potential of SHEDLOCK-2 as a next-generation CAR T platform that combines enhanced efficacy with a favorable safety profile.

## Discussion

Soluble BCMA shedding has emerged as a principal barrier to durable BCMA-directed CAR T cell therapy, and higher baseline concentrations of soluble BCMA are associated with poor clinical responses and early relapse^13–15^. Because BCMA shedding is enzyme dependent, small-molecule γ-secretase inhibitors such as nirogacestat can raise surface BCMA more than 100-fold and enhance CAR T cell cytotoxicity in preclinical and early clinical studies ^10, 12^. Systemic use, however, improves antigen stability at the cost of gastrointestinal and hematological toxicity that limits prolonged dosing^16^, arguing for confining γ-secretase inhibition to the tumor rather than delivering it systemically. The SHEDLOCK platform achieves this by encoding the inhibitor within the CAR T cell itself. SHEDLOCK-1 cells secrete the engineered peptide P5 locally, restricting BCMA cleavage to the tumor microenvironment while sparing non-target tissues. This design preserved membrane BCMA on myeloma cells, reduced soluble BCMA, enhanced cytokine production, and maintained robust cytotoxicity in vitro and in vivo, establishing local γ-secretase inhibition as a viable route to greater antigen stability without systemic exposure.

Preventing antigen loss alone proved insufficient, because chronic antigen exposure continued to drive T cell exhaustion and senescence. We therefore developed SHEDLOCK-2 cells secreting nxP, a Notch-modulating peptide. Because persistent Notch activation promotes terminal differentiation and suppresses telomerase, shortening T cell lifespan^26–28^, nxP was designed to interrupt this process; it restored telomerase activity, lowered p21, and promoted mitochondrial oxidative phosphorylation. The resulting cells acquired a memory-like phenotype with improved metabolic stability, reduced exhaustion markers, and enhanced longevity. Further, transcriptomic profiling confirmed enrichment of FOXO and oxidative phosphorylation associated with increased persistence of SHEDLOCK-2 cells.

To define how γ-secretase inhibition preserves telomeres, we dissected the Notch signaling pathway. Physiologically, γ-secretase cleaves Notch at the S3 site to release the Notch intracellular domain (NICD), which, together with RBPJ and MAML1, activates the Hes and Hey basic helix-loop-helix repressors that silence target genes^46–49^. Reasoning that telomerase might lie downstream of this output, we examined the hTERT promoter and identified three high-scoring Hes1 binding sites within the 2 kb region upstream of the transcription start site, at approximately −579, −282 and −163 bp. Two of these correspond to Class C (CACGCG) elements, a site class with established precedent for Hes1 occupancy ^50^, as (Kip1) promoter has been reported previously and mutation of the Class C site abolished Hes1-dependent repression Hes1 binding to Class C sites at the p27 ^50^. We confirmed Hes1 occupancy by chromatin immunoprecipitation in NTP- CAR T cells under tumor rechallenge, which was reduced in SHEDLOCK-2 cells as nxP abolished NICD generation, and silencing of Hes1 de-repressed hTERT and restored telomerase activity, identifying hTERT as a Notch target that governs T cell longevity. This axis explains how chronic antigen exposure undermines CAR T cell fitness, whereby sustained Notch signaling elevates Hes1, suppresses telomerase, and accelerates senescence and p21 accumulation. The underlying link between telomere dysfunction and T cell senescence, mediated through the ATM, CHK2, p53, and p21 cascade, is well-established^51–53^. Critically, the benefit of nxP derives from reduced Notch transcriptional output rather than from broad γ-secretase inhibition. Importantly, to deconvolute tumor-extrinsic (mBCMA retention) from cell-intrinsic (Notch-Hes1-hTERT) effects, we compared SHEDLOCK-2 and NTP-CAR T cells against cleavage-resistant BCMA-28tm targets, where shedding is already abrogated. Both groups showed comparable cytotoxicity and no differential telomerase or telomere preservation, confirming that the longevity benefit of nxP is cell-autonomous and Notch-dependent rather than antigen-retention-driven. Future studies using paired shedding-competent and shedding-resistant tumor models across extended in vivo time courses will be needed to fully quantify the relative weight of each mechanism.

A convergent metabolic mechanism reinforces this transcriptional suppression. Attenuation of Notch and Hes1 signaling shifts metabolism toward oxidative phosphorylation, upregulating FOXO and OXPHOS genes while downregulating glycolytic genes. The accompanying decrease in mitochondrial reactive oxygen species protects guanine-rich telomeric repeats from oxidative erosion, a recognised route to telomere attrition in chronically activated T cells ^53, 54^ . Together, telomerase-dependent synthesis and reduced oxidative loss act in concert to preserve telomere length, lower p21, and sustain the memory-like differentiation and durable persistence that distinguish SHEDLOCK-2 from conventional CAR T cells under repeated challenge. These findings are consistent with the well-established role of telomere shortening and senescence in driving CAR T cell functional decline ^52–54^. They also connect to a broader effort to extend CAR T cell durability through metabolic and signaling engineering. Secreted-payload designs, such as a fifth-generation BCMA CAR co-expressing an anti-PD-L1 fragment or a B-NOXA apoptosis-directing fusion, act through checkpoint inhibition or targeted killing, whereas metabolic strategies, including PGC-1α activation, mTORC1 regulation, IL-15 or AKT3 modulation, and our previously described PROTAC-mediated AKT3 degradation that raises FOXO4, enhance oxidative metabolism and persistence^29, 33, 56–60, 62^ . Telomerase-directed T cells and transcriptional-enhancer CARs that boost BCMA and mitochondrial fitness likewise reduce exhaustion, and we have previously shown that GLP-1-engineered CAR T cells improve mitochondrial output and delay senescence ^42, 55, 60, 61^. SHEDLOCK-2 is distinguished from these approaches by integrating antigen stabilisation with metabolic reprogramming in a single, locally acting construct.

By coupling antigen stabilisation to metabolic reprogramming, SHEDLOCK-2 cells sustained BCMA engagement, oxidative metabolism, and a memory-like phenotype, translating into durable tumor control even after repeated antigen challenge; unlike checkpoint-only designs, they improve both antigen availability and T cell longevity within one construct, and the platform remains extensible to metabolic enhancers such as PGC-1α activators and to apoptosis-directing modules such as the B-NOXA fusion ^57, 63^. This is achieved without the off-tumor toxicity of systemic γ-secretase inhibition, because peptide release is spatially restricted. Custom sandwich ELISAs confirmed that circulating P5 and nxP peaked early after infusion and declined below the limit of detection by day 45, whereas bone-marrow concentrations at sacrifice exceeded contemporaneous blood levels by several-fold. This compartmentalisation confirms preferential peptide accumulation within the tumor microenvironment, where local concentrations were sufficient for sustained soluble-BCMA suppression and membrane-BCMA rescue, while systemic exposure waned in parallel with CAR T cell contraction. Consistent with this compartmentalisation, 90-day monitoring of mice given SHEDLOCK-2 cells showed no on-pathway, biochemical, or organ-related toxicity. nxP also proved sequence specific and target restricted in vitro, with no off-target activity. Viability was unchanged across epithelial, B, T, NK, and neuronal cells, and a scrambled-sequence peptide neither suppressed soluble BCMA nor altered cell viability. A dose-response screen at concentrations far above those measured in the bone marrow showed no cytotoxicity, and the peptide was inactive against a cleavage-resistant BCMA mutant, a stringent on-target control. Together, these findings suggest that residual on-pathway Notch modulation may be the main off-target effect warranting clinical monitoring.

Besides defining the mechanism, our preclinical findings identify CAR T-intrinsic correlates that may inform future clinical development. Profiling the product before infusion and tracking it under tumor-rechallenge conditions revealed features that correlated with durable tumor control. Telomere length and telomerase activity, measured before infusion and under tumor rechallenge, correlated with CAR T cell persistence and survival^24^. Uniquely, Hes1 transcript levels, which were several-fold lower in SHEDLOCK-2 cells and correlated inversely with telomere length but positively with p21, provide a direct pharmacodynamic readout of nxP target engagement. Combined with soluble BCMA and with senescence, exhaustion, and memory markers, this telomere, metabolic, and Notch panel offers a coherent, multi-parameter framework for correlative endpoints and exploratory release criteria in first-in-human evaluation, though prospective clinical validation will be required before any parameter can be designated a predictive biomarker.

Finally, the utility of this platform extends well beyond BCMA. Because the secreted-peptide module acts independently of the antigen-recognition domain, it can be transferred to other targets and to dual-antigen constructs, including those directed against GPRC5D, TACI, CD70, and CD19^64, 65^, carrying localized γ-secretase inhibition and metabolic reprogramming into any CAR architecture and building on the bispecific and dual-antigen strategies established by several groups ^66–68^. In doing so, SHEDLOCK addresses the two principal mechanisms that most often drive relapse, antigen escape and T cell exhaustion, within a single, modular design. This dual action positions SHEDLOCK not as an incremental modification but as a broadly adaptable framework for the next generation of durable T cell therapies, with the potential to improve outcomes across multiple hematological and solid malignancies.

### Translational considerations and limitations

Several factors will influence clinical translation. The combined transgene approaches the lentiviral packaging limit and may modestly lower titre, so GMP-grade production will require titre optimisation and defined release criteria for the secreted-peptide module. Immunogenicity is a further consideration, and although nxP and P5 were designed for low predicted MHC binding, anti-drug antibody (ADA) monitoring and in vitro human T-cell activation assays (e.g., ELISpot-based immunogenicity screening) will be essential components of the Phase I clinical protocol, as in silico MHC-binding prediction alone cannot fully exclude immunogenic risk. SHEDLOCK addresses antigen shedding and exhaustion but cannot alone prevent antigen escape through biallelic BCMA loss or splicing, which supports combination with dual-antigen designs such as BCMA with GPRC5D or CD19. Acquired resistance to nxP or P5 is unlikely given the essentiality of γ-secretase, though substrate processing and Notch activity will still be monitored as correlative measures. Overall, SHEDLOCK is best viewed as a modular enhancement that complements existing BCMA CAR T backbones rather than replacing them.

## EXPERIMENTAL MODEL AND STUDY PARTICIPANT DETAILS

### Cell lines

The cell lines MM.1R, HEK293 T, HS-5, K562, HeLa, HepG2, A549, SH-SY5Y, Raji, NALM-6, and Jurkat were obtained from ATCC, while MM.1S was purchased from BPS Biosciences. All cell lines were cultured according to the manufacturer’s recommendations. HEK293 T cells were maintained in DMEM supplemented with 10% fetal bovine serum (FBS) and were passaged upon reaching 70-80% confluency. MM.1R, MM.1S, Jurkat, and Raji cells were cultured in RPMI-1640 medium supplemented with 10% FBS and sub-cultured at 70-80% confluency. HS-5 cells were maintained in DMEM with 10% FBS and seeded at a density of 5, 000 cells/cm². K562 cells were cultured in IMDM supplemented with 10% FBS. HeLa and HepG2 cells were maintained in EMEM supplemented with 10% FBS, while SH-SY5Y cells were cultured in a 1:1 mixture of EMEM and Ham’s F-12 K nutrient medium containing 10% FBS. Cell lines were cultured at 37**°**C, 5% CO2, and 95% humidity. All cell lines were routinely tested for mycoplasma contamination.

### Human Samples

Peripheral blood mononuclear cells (PBMCs) and whole blood were obtained from healthy adult donors (25-60 years) at Apollo Indraprastha Hospital after written informed consent, as described in our previous study^42^. All procedures complied with institutional ethical and regulatory standards and were approved by the Apollo Hospitals Educational and Research Foundation, Institutional Ethics Committee (IRB Ethics Waiver, Application No. IAH-BMR-064/07-23). Patient PBMCs were collected from 45 patients following written informed consent^42^. In addition, bone marrow aspirates from patients with multiple myeloma (ages 30-60 years; 22 males and 15 females) were obtained from Sarvodaya Hospital, Faridabad, Haryana, India, after written informed consent and in accordance with institutional ethical policies and regulatory requirements for research and development (SHRC-IEC/2026/51). All blood and bone marrow samples used in this study were surplus specimens remaining after routine diagnostic marker analyses and would otherwise have been discarded.

### Animal Experiments

NOD.Cg-Prkdc^scid^ Il2rg^tm1Wjl^/SzJ (NSG, Stock no.#005557) and NOD-Prkdc^em26Cd52^Il2rg^em26Cd^^22^/NjuCrl (NCG) (Strain Code: 572) mice (both male and female), aged 6 to 8 weeks, were obtained from The Jackson Laboratory and Charles River Laboratories, respectively, and maintained under specific pathogen-free conditions in individually ventilated cages at 22°C with a 12-h light/dark cycle and access to food and water. These strains were used for all multiple myeloma xenograft studies, SHEDLOCK-1 and SHEDLOCK-2 efficacy assessments, serial-transfer experiments, and the patient-derived xenograft (PDX-MM) model. Tumor engraftment, CAR T cell infusion timing, and longitudinal monitoring of bioluminescence, circulating human Ig, CAR T cell persistence, and bone-marrow infiltration were performed exactly as described and follow the methodological framework previously reported for NSG-based CAR T cell studies^42^.For cytokine release syndrome (CRS) modelling, 4- to 6-week-old female SCID-beige mice (C.B-Igh-1b/GbmsTac-Prkdc^scid^ Lyst^bg-N7^) were obtained from Taconic Biosciences and housed under identical environmental conditions. These mice were selected based on their heightened sensitivity to T cell derived cytokine responses and were used to evaluate acute inflammatory toxicity associated with SHEDLOCK CAR T cell infusion. Serum cytokines, liver and renal enzyme panels, and tocilizumab mediated rescue experiments were conducted as outlined in the manuscript and consistent with previously validated CRS models. All procedures, including tumor inoculation, CAR T cell administration, blood and bone-marrow harvests, analgesia, and monitoring, were performed in accordance with institutional guidelines. The animal experiments were conducted following approval from the respective Ethics Committee for Animal Experiments. All animal experiments were approved by the Institutional Animal Ethics Committee of Meril Diagnostics, Vapi, Gujarat India and CSIR-Institute of Chemical Biology, Kolkata, West Bengal, India.

## STAR METHODS

### Computational design of γ-secretase-modulatory peptides

The peptide design pipeline was made using RFDiffusion v1.1^69, 70^ and AlphaFold2-Multimer v2.3.2^71^. Publicly available pre-trained weights were used; no fine-tuning was performed. RFDiffusion has been trained by its developers on the protein-structure subset of the PDB (approximately 25, 000 chains at <30% sequence identity), and ProteinMPNN was trained on a complementary curated PDB set; full training datasets are documented in the original publications. No new training data were generated or curated for this study.

The full-length human BCMA monomer (UniProt Q02223; AlphaFold2 model AF-Q02223-F1) served as the immobile target. Residues 55-77, spanning the juxtamembrane and transmembrane segment containing the experimentally mapped γ-secretase cleavage site^11^, were defined as the hotspot. The target was positioned in an implicit POPC: DOPC bilayer plane during RFDiffusion sampling using a contig specification permitting linear binders of 15-25 residues. RFDiffusion was run with 50 denoising timesteps (T = 50), translation noise scale 0.5 and rotation noise scale 1.0; 500 backbones were generated per independent run, across three independent runs, yielding 1, 500 backbones in total. For each backbone, ProteinMPNN designed 5 sequences at sampling temperature 0.1 with the target chain held fixed, producing 7, 500 candidate binder sequences. Each binder-BCMA complex was then re-folded with AlphaFold2-Multimer (5 models per complex), and the top 200 candidates were carried forward based on interface pLDDT.

Candidates were filtered using the following hard cutoffs applied sequentially: (i) interface pLDDT > 80; (ii) predicted aligned error at the binder-target interface (pae_interaction) < 10 Å; (iii) Cα-RMSD of the binder to its design model < 2 Å; (iv) ipTM > 0.65; and (v) Kyte-Doolittle hydrophobicity index between 0.4 and 0.6 to balance membrane insertion against aggregation risk. These filters reduced the 7, 500 designs to 64 high-confidence candidates (**Table 1**); the top five (P1-P5) were selected for synthesis using a composite score defined as (interface pLDDT x ipTM) / pae_interaction, with hydrophobicity used as a tiebreaker.

The naturally derived nxP peptide series was generated through a parallel inverse-folding pipeline distinct from the de novo RFDiffusion workflow used for P1-P5 above. Starting templates were the Notch1 negative-regulatory region (PDB 3I08; residues 1664-1737) and the APP transmembrane domain (PDB 2LP1; residues 686-726), both well-characterised native γ- secretase substrates. Each template was positioned onto the cryo-EM structure of human γ-secretase (PDB 5A63), with the catalytic core of presenilin-1 (PS-1); specifically, the aspartate dyad (D257 and D385) and the adjacent substrate-docking residues lining the intramembrane catalytic groove, which is defined as the fixed interface. ProteinMPNN was then run in inverse-folding mode at a sampling temperature of 0.05, generating 100 candidate sequences per template while preserving the backbone geometry of the substrate helix. Candidates were re-folded with AlphaFold2-Multimer for interface validation and subsequently subjected to 100-ns all-atom molecular-dynamics simulations in a POPC: DOPC bilayer, followed by MM/GBSA binding free-energy estimation against PS-1. Substitutions that improved predicted binding affinity (ΔΔG ≤ -2 kcal mol^-1^) without compromising helical integrity (Cα-RMSD < 1.5 Å) or introducing steric clashes within the catalytic groove were retained, yielding two lead candidates; nx-P1 (APP-derived) and nx-P2 (Notch1-derived). nx-P2 was prioritised for downstream functional characterisation, and hereafter designated nxP, on the basis of its higher PS-1 occupancy, lower interfacial RMSF, and persistent contact with the catalytic aspartates across three independent simulation replicates.

### Molecular docking and binding free energy estimation

Peptide-BCMA interactions were evaluated using a structure-guided assembly and binding free-energy estimation approach consistent with the molecular-dynamics framework described above. Structural models of peptide-BCMA TM complexes were generated by positioning candidate α-helical peptides parallel to the BCMA TM helix within a membrane context, guided by steric complementarity and predicted helix-helix interfaces. Binding energetics and interface stability were assessed using MM/GBSA calculations implemented within the AMBER22 workflow. Free-energy estimates (ΔG_bind) were computed from equilibrated MD trajectories, and residue-level energy decomposition was performed to identify key contributions from hydrophobic packing, hydrogen bonding, and electrostatic interactions at the peptide-BCMA interface. Peptides were prioritized based on predicted binding free energy, persistence of interfacial contacts across simulation trajectories, and stable orientation along the BCMA TM axis proximal to the γ-secretase cleavage region. These analyses enabled identification of peptide candidates predicted to form sustained, cleavage-blocking interactions with BCMA, supporting selection of P5 for downstream experimental validation.

### Molecular dynamics simulation in lipid bilayer systems

All-atom MD simulations were performed using AMBER22 to evaluate the stability of peptide-BCMA interactions within a membrane environment. Peptide-BCMA TM complexes were embedded in a mixed POPC: DOPC lipid bilayer, solvated with explicit water molecules, and neutralized with counter-ions to achieve physiological ionic strength. System setup, membrane embedding, and parameterization were performed following standard AMBER membrane simulation protocols. Each system underwent energy minimization to remove steric clashes, followed by stepwise equilibration under constant volume (NVT) and constant pressure (NPT) conditions prior to production runs. Production MD simulations were carried out for 200 ns (unless mentioned), with three independent replicates per peptide-BCMA complex, under periodic boundary conditions at physiological temperature. Trajectory analyses were performed to assess structural stability and peptide engagement, including root mean square deviation (RMSD), peptide-TM contact frequency, hydrogen-bond persistence, helix orientation relative to the BCMA TM axis, and residue-level interaction profiles. Per-residue energy decomposition and time-resolved contact analyses were used to evaluate persistence of peptide interactions near the BCMA juxtamembrane γ-secretase cleavage region.

### Structural modelling and visualization of BCMA peptide interactions

Structural models of BCMA-peptide complexes generated from docking and MD simulations were visualized using PyMOL and ChimeraX to assess residue-level interactions and overall spatial orientation at the γ-secretase cleavage interface. Electrostatic surface maps were computed using the APBS plugin to evaluate charge complementarity and identify regions of favorable peptide anchoring along the BCMA juxtamembrane helix. Representative structural snapshots from equilibrated MD trajectories were examined to determine solvent-exposed and membrane-facing residues, focusing on motifs predicted to sterically occlude γ-secretase access. These visualization analyses aided in refining peptide candidates by highlighting stable contact regions, helical packing behavior, and structural features supporting cleavage-site obstruction.

### CAR construct design and AI-guided humanization

CAR constructs used in this study, including the AI-humanized BCMA CAR (HmzBCMA), were designed using a computational pipeline integrating AI-guided ScFv humanization, adapted from our previously established workflow^72^. Murine anti-BCMA scFv sequences (clone: 4C8A) were retrieved from published literature and subjected to an AI-based humanization framework incorporating machine-learning-driven CDR grafting, de-immunization scoring, and iterative framework refinement to maximize humanness while preserving antigen-binding residues. Humanization of the murine 4C8A clone was performed using BioPhi (v1.0). The AI model evaluates structural compatibility, CDR loop conformation, solvent exposure, and potential immunogenic liabilities prior to generating candidate humanized variants. Top-ranked humanized BCMA scFvs were structurally modelled using AlphaFold2 (DeepMind) for three-dimensional structure prediction and subsequently refined in UCSF Chimera (v1.16) with MODELLER to preserve native CDR geometry and paratope integrity. These models were docked for antigen binding with human BCMA Antigen using the HADDOCK webserver(v2.4).To ensure structural stability and retention of BCMA-binding interfaces, the modelled scFvs underwent molecular-dynamics (MD)-based structural validation using AMBER22, assessing framework packing, CDR flexibility, and paratope stability over simulation trajectories. Variants demonstrating optimal structural stability and conserved antigen-binding features were selected for downstream CAR engineering.

### CAR lentiviral vectors

All the CAR plasmids which are used in this study were prepared in third generation self-inactivating lentiviral (LV) backbone using synthetic gene fragments that were designed in house and obtained from GenScript, and the cloning and lentivirus production steps followed the strategy described by us in our previous study ^42^. The BCMA CAR plasmid was made by scFv derived from the murine 4C8A clone together with a CD8 hinge and a CD8 transmembrane domain, followed by CD28, 4-1BB co-stimulatory domains and the CD3ζ signaling domain, and the AI humanized scFv (HmzBCMA CAR) was cloned in exactly the same arrangement so that the two constructs differ only in the antigen binding region, while the CD19 CAR (FMC63 clone), which was used as an antigen independent control for peptide secretion, was also cloned in this same configuration. Further, 4-1BB based BCMA targeting murine derived scFv used in the ide-cel type construct was obtained from the patent (Patent WO2021091978A1) and cloned in the same vector.

To obtain the peptide secreting SHEDLOCK constructs, synthetic fragments encoding the γ-secretase modulatory peptide P5 (SHEDLOCK-1), the modified Notch derived peptide nx-P2, (SHEDLOCK-2), or a non-targeting peptide of comparable length (NTP, used as control) were obtained from GenScript, each carrying an IL-2 signal peptide at the 5′ end to direct secretion of the peptide and joined to the CAR open reading frame through a self-cleaving T2A element, and these fragments were cloned in frame into the HmzBCMA CAR plasmid to generate single plasmid vectors in which the CAR and the secreted peptide are expressed from one promoter as a single transcript; the CD19-P5 and CD19-NTP plasmids were prepared in the same manner by replacing the BCMA scFv with the FMC63 scFv. The other plasmids were prepared in the same LV backbone, and these included the cleavage resistant BCMA variant (BCMA-28tm), and the wild type BCMA construct that was used to generate the stable HEK-BCMA and K562-BCMA cell lines, both cloned from synthetic fragments; full length Hes1 cloned downstream of the EF1α promoter (EF1α-pHes1) for overexpression studies; the constitutively nuclear and γ secretase independent Notch1 intracellular domain (NICD1-ΔE), which was obtained as a synthetic gene fragment and cloned into the LV expression plasmid; short hairpin RNA (shRNA) constructs against Notch1, which were designed using the shRNA Design Tool (IDT) and cloned as described by us earlier, with a scrambled control ^42^. The hTERT proximal promoter firefly luciferase reporter and its Hes1 site mutant, which were maintained in pGL3-Basic were cloned into pLenti and the mCherry and firefly luciferase fragments that were cloned into the same LV backbone to generate the reporter tumor lines MM.1S, MM.1R, NALM6 and Raji ^42^.

For lentivirus production, HEK293 T cells were seeded in T-175 flasks at a density of 4.2×10^6^ cells in 30 mL DMEM containing 10% fetal bovine serum, and after 48 hours cells were co-transfected using PEI (Sigma) or PEIpro (Polyplus) with the third generation packaging plasmids pRSV-Rev, pRRE and pMD2.G (VSV-G) along with the respective CAR or accessory transgene plasmid. The cells were kept in the transfection medium for 48 h, after which the medium was changed and the cells were cultured for a further 24 h. Viral supernatants were collected at 48 h and 72 h after transfection, clarified, concentrated using TFF and chromatography systems as described previously ^72^ or Lenti-X concentrator (Takara), and the resulting pellet was resuspended in OptiMEM and stored as single use aliquots at -80 °C ^72^. The functional titre of every batch was determined by serial dilution of the concentrated LV particles transduced in HEK293 T cells followed by measurement of transduction efficiency by flow cytometry, and titre matched preparations were used to transduce activated primary T cells at a multiplicity of infection 5 (MOI 5), so that the level of CAR surface expression was comparable between the NTP, SHEDLOCK-1 and SHEDLOCK-2 groups in all comparative experiments. The complete list of plasmids, synthetic gene fragments and shRNA sequences is provided in the key resource table. To compare CAR surface expression across constructs, primary human T cells from independent donors were transduced in parallel with empty vector, Hmz-BCMA CAR, SHEDLOCK-1 or SHEDLOCK-2 lentiviral vectors at MOI 5 and analysed on day 7 post-transduction. CAR surface expression was detected by staining with a BCMA CAR detection antibody (Miltenyi Biotec) followed by streptavidin-FITC, and the percentage of CAR-positive cells was quantified by flow cytometry (BD FACSLyric). For lentiviral titre determination, HEK293 T cells were transduced with serial volumes (0, 0.25, 0.5, 1, 2 and 3 μL) of concentrated lentiviral supernatant harvested at 48 h from producer cells. After 72 h, CAR-FITC expression was assessed by flow cytometry, and the infectious titre (transducing units per mL; TU/mL) was calculated from the linear range of the dose-response curve using the formula: TU/mL = (% CAR-positive cells × cell number at transduction) / (volume of virus in mL).

The detailed information about the vector sequence and the design along with the vector maps is provided in **Table 5**.

### CAR T cell generation, transduction, and expansion

Peripheral blood mononuclear cells (PBMCs) were isolated from whole blood obtained from healthy donors using Ficoll-Paque Plus based density-gradient centrifugation. T cells were enriched and activated by positive selection with CD3/CD28 Dynabeads, with bead-only stimulation. Activated T cells were cultured in AIM-V serum-free medium supplemented with 10% FBS and recombinant human IL-2 (100 IU/mL). On day 0 (D0), T cell purity was assessed by multiparametric flow cytometry using lineage and exclusion markers, including CD3, CD4, CD8, CD16, CD14, and CD56, to confirm efficient depletion of monocytes, NK cells, and other non-T cell contaminating populations prior to lentiviral transduction. All the antibodies are in different channels and FMOs were generated while flow cytometry acquisition wherever needed.

On Day 1 (D1), 1 × 10^6^ activated T cells were transduced with lentiviral vectors encoding the respective BCMA CAR constructs at MOI 5 in complete AIM-V medium. Cells were maintained in culture until Day 5 (D5), when magnetic de-beading was performed to remove CD3/CD28 beads. Immediately following bead removal, CAR T cells were transferred to fresh AIM-V medium supplemented with IL-7 and IL-15 (84 IU and 100 IU respectively) and expanded until Day 7 (D7), with media replenished every 48 hours. A seeding density of 1 × 10^6^ cells/mL was maintained throughout the expansion phase to support optimal proliferation and viability. On Day 7, CAR T cells were harvested and stained with the BCMA CAR Detection antibody to confirm surface CAR expression. Flow cytometric acquisition was performed and data was analysed using FlowJo software FlowJo 10.10.0 to quantify CAR positivity and T cell purity.

### In vitro BCMA shedding assay

The peptides were synthesized by GenScript Biotech (Singapore). Peptides were reconstituted in 1X PBS, and applied to MM.1S cells at various concentrations. For experiments involving peptide-secreting CAR constructs, CAR T cells were expanded in AIM-V serum-free medium supplemented with 10% fetal bovine serum and recombinant human IL-2 (100 IU). On day 5 of expansion, cultures were transferred to fresh AIM-V medium containing IL-2 and seeded at 1 × 10^6^ cells/mL, followed by overnight incubation. The following day, CAR T cell cultures were centrifuged at 300 rcf for 10 min, and the resulting cell-free supernatant containing secreted peptide (secretory soup/supernatant) was collected and used directly for BCMA shedding assays, while the corresponding cell pellets were retained for downstream membrane BCMA analysis. Peptide and secretory-soup treatments were performed in serum-free RPMI medium. Following treatment, plates were centrifuged at 1, 200 rpm for 5 min, and clarified supernatants were collected and stored at -80°C until batch analysis. Soluble BCMA was quantified using the human BCMA/TNFRSF17 Duo Set ELISA (Cat# DY193, R&D Systems) according to the manufacturer’s instructions, with recombinant BCMA-Fc used to generate standard curves. In parallel, the γ-secretase inhibitor DAPT was used as a positive control for shedding inhibition.

Corresponding cell pellets from each condition were processed for membrane BCMA flow-cytometry analysis to correlate soluble BCMA reduction with surface BCMA retention. Absorbance for soluble BCMA was measured using a VICTOR Nivo multimode microplate reader (PerkinElmer), and data were analyzed using GraphPad Prism software.

### Custom sandwich ELISA for detection of P5 and nxP peptides

Blood and bone marrow samples were collected from mice at the indicated time points for quantification of the secreted peptides P5 and nxP. Peptide concentrations were measured using two custom sandwich ELISAs built around target-derived capture reagents. For P5, a recombinant BCMA transmembrane peptide (rBCMA-TM) served as the capture reagent; for nxP, a recombinant presenilin-1 substrate-docking peptide (rPS1) was used. Each reagent binds its cognate peptide through the same interface that mediates target engagement in vivo. Ninety-six-well microplates were coated overnight at 4 °C with the respective capture peptide (2 μg/mL in carbonate buffer, pH 9.6, 100 μL per well), washed three times with PBS containing 0.05% Tween-20 (PBST), and blocked with 3% BSA in PBS for 2 h at room temperature. Plasma and bone marrow supernatant samples were clarified by centrifugation, diluted in blocking buffer, and added to the wells (100 μL per well) alongside a standard curve of synthetic P5 or nxP peptide spanning 0.5 pg/mL to 100 ng/mL, prepared by serial dilution in assay buffer. Plates were incubated for 2 h at room temperature to allow capture of the peptide by the immobilized reagent. After washing, bound peptide was detected with HRP-conjugated rBCMA-TM or rPS1 (1 μg/mL, 1 h at room temperature). Wells were developed with TMB substrate, the reaction was stopped with 2 N H₂SO₄, and absorbance was read at 450 nm with 570 nm reference correction on a microplate reader. Peptide concentrations were interpolated from the standard curve using a four-parameter logistic fit. The lower limit of detection (LLOD) was 0.5 pg/mL, defined as the concentration yielding a signal three standard deviations above the mean blank, and the lower limit of quantification (LLOQ) was 1 pg/mL. Intra-assay coefficients of variation were below 10% across the quantifiable range. Samples falling below the LLOD were assigned a value of zero for graphing and statistical purposes. Assay specificity was confirmed using non-targeting peptide (NTP) controls, which yielded signals at background level.

### In silico identification of Hes1 binding sites

The proximal hTERT promoter was analyzed to identify candidate recognition motifs (class C sites, core sequence CACNAG, and N-box elements) using the JASPAR position weight matrix for Hes1 together with a manual search for canonical E-box/N-box sequences. Three high-scoring sites located near -579, -282, and -163 bp were selected for experimental validation.Hes1 binding elements. A comprehensive in silico analysis was performed to predict Hes1 transcription factors binding site prediction within the 2000-bp region upstream of the hTERT transcription start site . Using all three available JASPAR Hes1 position-weight matrices (MA1099.1, MA1099.2 and MA1099.3) 13 high confidence binding sites were identified. The 2 kb genomic region upstream of the hTERT transcription start site was retrieved from GRCh38/hg38 reference genome using the TSS of the canonical TERT-201 transcript (ENST00000310581). Because hTERT is located on the minus strand, the promoter sequence was reverse complemented and screened for consensus Hes1

### Chromatin immunoprecipitation (ChIP)-qPCR

CAR T cells (NTP, SHEDLOCK-2) were expanded and subjected to the in vitro tumor re-challenge protocol before harvest. Approximately 1 to 2 × 10^7^ cells per condition were cross-linked with 1% formaldehyde for 10 min at room temperature, and cross-linking was quenched with 125 mM glycine for 5 min. Cells were washed in cold PBS, lysed, and chromatin was sheared to fragments of 200 to 500 bp by sonication. Solubilized chromatin was precleared and incubated overnight at 4 °C with an anti-Hes1 antibody or a matched isotype IgG control, and immune complexes were captured on protein A/G magnetic beads. After sequential washes, chromatin was eluted, cross-links were reversed at 65 °C overnight, and DNA was purified following proteinase K and RNase treatment. Enrichment at the three predicted Hes1 sites in the hTERT promoter was quantified by qPCR using site-specific primers, and results were expressed as percentage of input, with a control genomic region serving as a negative locus. Hes1 occupancy was compared between NTP-CAR T cells and SHEDLOCK-2 cells.

### hTERT proximal promoter-luciferase reporter assay

The proximal hTERT promoter region containing the three predicted Hes1 sites was cloned upstream of the firefly luciferase gene in a pGL3-Basic reporter vector to generate the wild-type reporter. A mutant reporter was produced by site-directed mutagenesis of all three Hes1 binding sites to disrupt the core recognition motifs. Reporter constructs were co-transfected with a Hes1 expression plasmid or empty vector into the target cells, together with a Renilla luciferase plasmid as an internal transfection control. Cells were lysed 24 to 48 h after transfection, and firefly and Renilla activities were measured with a dual-luciferase assay on a luminometer. Firefly signal was normalized to Renilla, and promoter activity in the presence of Hes1 was expressed relative to the empty-vector control to quantify repression.

### Hes1 expression under in vitro tumor re-challenge (TR) conditions

CAR T cells were subjected to repeated antigen exposure in the in vitro TR model, and cells were collected at defined intervals. Hes1 expression was measured over time by quantitative RT-PCR and confirmed at the protein level by immunoblot or flow cytometry, with expression normalized to a housekeeping gene or loading control. Hes1 levels in NTP-CAR T cells and SHEDLOCK-2 cells were compared in parallel with telomerase activity, exhaustion markers, and CAR T persistence measured from the same cultures.

### Hes1 knockdown (siRNA) and overexpression

For transient knockdown, CAR T cells under TR conditions were transfected with Hes1-targeting small interfering RNA (siRNA) or a non-targeting control siRNA by electroporation, and knockdown efficiency was verified by RT-PCR and immunoblot. For gain-of-function experiments, resting CAR T cells were transduced with a lentiviral or plasmid vector encoding full-length Hes1 or an empty control vector. After knockdown or overexpression, telomerase activity, hTERT expression, exhaustion markers, and CAR T persistence were assessed as described. The siRNA sequences are provided in the source table.

### Notch1 knockdown (shRNA)

Notch1 was stably silenced using a lentiviral short hairpin RNA (shRNA) targeting Notch1, with a scrambled shRNA as control. The plasmid used for Notch1 shRNA cloning and the cloning procedure have been described previously^42^. Briefly, CAR T cells were transduced, selected where appropriate, and knockdown was confirmed by RT-PCR and immunoblot for Notch1 and NICD. Hes1 expression and telomerase activity were then measured after TR. For epistasis experiments, Notch1-knockdown CAR T cells were additionally transduced with the Hes1 overexpression vector, and telomerase activity, exhaustion, and durability were evaluated.

### NICD1-ΔE lentiviral construct and downstream reconstitution

To restore Notch transcriptional output independently of γ-secretase, a lentiviral construct encoding NICD1-ΔE, a constitutively nuclear, γ-secretase-independent form of the Notch1 intracellular domain, was generated. The NICD1-ΔE sequence was obtained commercially as a synthetic gene fragment and cloned into the lentiviral expression plasmid, which has been described previously ^42^. SHEDLOCK-2 cells were transduced with NICD1-ΔE or a control vector. Transduced cells were then evaluated for Hes1 expression and telomerase activity, and p21 expression, and results were compared between NICD1-ΔE or a control vector in SHEDLOCK-2 cells. To deconvolute the cell-autonomous Notch-dependent benefit of nxP from its effect on antigen retention, an in vitro tumor-rechallenge model was established using cleavage-resistant HEK-BCMA28^tm^ target cells alongside wild-type HEK-BCMA^wt^ controls. NTP-CAR T and SHEDLOCK-2 CAR T cells were co-cultured with either HEK-BCMAwt or HEK-BCMA28tm cells and rechallenged on days 9, 11, 13, 15, 17 and 19 as described for the standard TR assay. At the day-19 endpoint, CAR T cell survival was assessed by 7-AAD exclusion and flow cytometry. Telomere length was measured by quantitative PCR using the T/S ratio method, and telomerase activity was quantified by the real-time PCR-based TRAP assay (TRAPeze RT kit), both as described above. Comparisons between NTP-CAR T and SHEDLOCK-2 across the two target-cell conditions were used to determine whether the telomere-preserving benefit of nxP persists when BCMA shedding is already abolished.

### Membrane BCMA quantification

Following peptide or secretory-soup treatment, cells were harvested and washed twice with 1X PBS to remove residual media and peptide. Total viable cell numbers were recorded using a standard automated cell counter to ensure uniform staining conditions. For quantitative assessment of surface BCMA expression, approximately 0.5 × 10^6^ cells per condition were resuspended in BD staining buffer and stained with anti-BCMA/CD269 antibody together with 7-AAD for viability discrimination, following the manufacturer’s recommendations. Antibody master mixes were prepared fresh at the specified dilutions, and samples were incubated on ice for 30 min in the dark. After incubation, cells were washed twice with 1X PBS to remove unbound antibody and resuspended in 1X PBS for flow cytometry acquisition. Flow cytometric analysis was performed on a BD FACSLyric cytometer (BD Biosciences), acquiring at least 50, 000 events per sample. Data were analyzed using FlowJo software, employing standard gating strategies to exclude debris, doublets, and 7-AAD⁺ non-viable cells prior to quantifying BCMA/CD269 surface expression.

### Super-resolution microscopy and imaging analysis

Super-resolution imaging was performed to visualise CAR scFv surface expression on engineered T cells and membrane-bound BCMA (CD269) on MM.1S myeloma cells, following previously established protocols from our laboratory^72^.For CAR ScFv visualisation, briefly 1.5 × 10^6^ CAR T cells were harvested from actively expanding cultures, washed twice with 1X PBS, and stained on ice with the BCMA CAR Detection Antibody (Miltenyi Biotech) for 15 min. Cells were then incubated with streptavidin-PE for an additional 15 min on ice, with all steps performed in the dark to minimise fluorophore photobleaching and receptor internalisation. For membrane BCMA retention, MM.1S cells were processed in parallel by washing twice with 1X PBS and staining with anti-BCMA/CD269 antibody (according to the manufacturer’s protocol) to assess BCMA surface distribution. After staining, CAR T and MM.1S cells were washed thoroughly and fixed in 4% paraformaldehyde for 30 min at room temperature, followed by two PBS washes. Fixed cells were mounted onto poly-L-lysine–coated slides using Prolong Gold Antifade Mountant and allowed to cure overnight in the dark. Super-resolution imaging was carried out on a Zeiss ELYRA PS.1 system equipped with a 63×/1.40 NA oil-immersion objective (Plan-Apochromat 63×/1.40 Oil DIC) using Structured Illumination Microscopy (SIM). Images were acquired in 640 nm laser 150 mW or 488 nm 100 mW laser line, reconstructed using Zeiss Zen software with standard SIM parameters. Final analysis including fluorescence-intensity quantification, surface distribution mapping, and comparative visualisation of CAR scFv versus BCMA membrane topology was performed using Fiji (ImageJ), applying consistent thresholds and segmentation settings across all conditions.

### Cell viability and proliferation assays

Cell viability and proliferation assays were performed to assess (i) antigen-dependent CAR T cell fitness following co-culture with multiple myeloma targets and (ii) cellular safety across diverse cell types following peptide exposure, using CFSE-based proliferation assays and viability staining as described previously with few modifications^42^. For CAR T cell viability and proliferation, CAR T cells were harvested on Day 7 post-transduction and cultured exclusively in AIM-V CTS medium throughout the assay. Cells were washed twice with 1X PBS and labelled with Cell Trace™ CFSE (Thermo Fisher Scientific, Cat. No. C34554) according to the manufacturer’s instructions. Briefly, CAR T cells were resuspended at 1 × 10⁶ cells/mL in pre-warmed 1X PBS containing CFSE at a final concentration of 10 µM and incubated for 20 min at 37°C with intermittent mixing. Labelling was quenched by addition of AIM-V CTS medium, followed by two washes to remove excess dye. CFSE-labelled CAR T cells were rested for 30 min at 37°C prior to co-culture. CFSE-labelled CAR T cells were co-cultured with MM.1S or MM.1R multiple myeloma cells at defined effector-to-target (E: T) ratios as indicated in the figure legends, in AIM-V CTS medium. Co-cultures were maintained and at each time point, cells were harvested and stained with 7-aminoactinomycin D (7-AAD) to exclude non-viable cells. Flow-cytometric acquisition was performed on a BD FACSLyric cytometer, acquiring a minimum of 50, 000 events per sample. Data were analyzed using FlowJo software with standard gating strategies including lymphocyte gating, doublet exclusion, dead-cell exclusion, and quantification of CFSE dilution within the viable CAR T cell population. CAR T cell viability was expressed as the percentage of 7-AAD⁻ cells, while proliferation was quantified based on CFSE dilution and mean fluorescence intensity (MFI). For cellular safety assessment following peptide treatment, viability and proliferation assays were performed across multiple cell types representing epithelial, hematopoietic, neuronal, and immune lineages, as detailed in the Supplementary Information. Cells were cultured under lineage-appropriate conditions and treated with peptide (P5 or nxP) or vehicle control at the concentrations and durations specified in the figure legends. Following treatment, cell viability was assessed by 7-AAD staining, and proliferation was evaluated by CFSE dilution using Cell Trace™ CFSE. Flow-cytometric acquisition and analysis were performed as described above.

### Patient-derived MM sample processing, isolation and culturing

Primary bone marrow aspirates from multiple myeloma patients were obtained. Plasma cell enrichment was performed using a negative-selection strategy to recover untouched, viable myeloma cells directly from bone marrow. Briefly, 2 mL of fresh aspirate was incubated with RosetteSep Human Multiple Myeloma Cell Enrichment Cocktail at 50 µL/mL for 20-30 min at room temperature to allow crosslinking of unwanted leukocyte subsets to erythrocytes. The suspension was diluted with an equal volume of 1X DPBS and carefully layered over Lymphoprep density gradient medium without disturbing the interface, followed by centrifugation at 1, 200 x g for 20 min with the brakes off. Enriched myeloma cells were recovered from the interphase, washed, and immediately prepared for culture. As primary myeloma cells require stromal support to maintain viability and drug-response fidelity, cultures were established on HS-5 human bone marrow stromal cells that had been seeded 48 h earlier at a density of 1 × 10^5^ cells/mL in DMEM supplemented with 10% FBS to generate a uniform feeder monolayer, consistent with previously described surrogate bone-marrow microenvironment models^73^. Isolated myeloma cells were gently overlaid onto the stromal layer in RPMI medium supplemented with IL-6, BAFF, and APRIL, using concentrations validated for plasma cell survival^74, 75^. Recombinant human IL-6, BAFF, and APRIL/TNFSF13 were reconstituted in 1× PBS and added to cultures at final concentrations of 1 ng/mL, 200 ng/mL, and 200 ng/mL, respectively. Cells were maintained in RPMI medium supplemented with FBS at 37°C in a humidified incubator with 5% CO₂ and 95% humidity, with routine monitoring of viability by trypan blue exclusion and mycoplasma screening prior to downstream assays.

### In vitro immunogenicity assay

Immunogenicity of the murine and humanized BCMA scFvs was assessed using an ex vivo dendritic cell-PBMC co-culture assay, following the workflow described. Briefly, monocyte-derived dendritic cells (DCs) were generated from healthy-donor PBMCs by adherence enrichment and culture in RPMI supplemented with IL-4 and GM-CSF for DC differentiation. On maturation, DCs were pulsed with purified recombinant scFv proteins corresponding to the murine and humanized scFv. Antigen-loaded DCs were then co-cultured with autologous PBMCs at a 1:1 DC:PBMC ratio in complete RPMI medium (10% FBS) for 48 hours. Cytokine secretion from PBMCs specifically IFN-γ, IL-2, and IL-4 was measured in culture supernatants using ELISA Assay kits, as these cytokines represent key indicators of T cell alloreactivity.

### Cytokine and T cell functional profiling

Granzyme B expression in CAR T cells was quantified by intracellular flow cytometry. CAR T cells were co-cultured with MM target cells, treatment in In vivo mice models or in the peptide treatment samples in the presence of Brefeldin A to block cytokine secretion, followed by surface staining and fixation/permeabilization using BD Cytofix /Cytoperm. Intracellular Granzyme B was detected using anti-Granzyme B, and data were acquired under uniform cytometer settings. Granzyme B levels were reported as median fluorescence intensity (MFI) in viable CD3⁺ CAR T cells, with FMO controls used to establish gating boundaries. Secreted cytokines were measured from clarified culture supernatants (50 µl) using R&D Systems Quantikine ELISA kits: IFN-γ, IL-2, IL-4, IL-6, TNF-α and IL-1β . Samples were centrifuged at 300 × g for 5 min, stored at -80 °C, thawed on ice, and assayed in technical triplicates using kit-provided standards and controls. Plates were developed and read at 450 nm with a 570 nm reference for OD correction in Victor Nivo Multiplate reader. Cytokine concentrations were interpolated from standard curves and corrected for dilution, and final values are reported in pg/mL.

### Bio-layer interferometry (BLI) assay

Kinetic binding analyses between BCMA Antigen and the designed ScFvs were conducted using the Gator Biolayer Interferometry (BLI) system (Gator Bio Inc., USA) at room temperature. Biotinylated recombinant human BCMA-Fc (CD269) protein was immobilized onto streptavidin biosensors (Gator Bio, Cat. No. 160002). Prior to use, sensors were equilibrated in assay buffer [1× PBS supplemented with 0.02% (v/v) Tween-20] for 10Omin. For ligand loading, sensors were incubated in BCMA ScFv proteins (5–10Oµg/mL) until a loading signal of approximately 0.8-1.2Onm was achieved, followed by baseline equilibration in buffer. Analyte proteins (mu- ScFv and Hmz ScFv) were prepared in assay buffer at serial concentrations of 200, 100, 50, 25, 12.5, and 6.25OnM for kinetic evaluation. Association was monitored for 120Os, followed by dissociation in assay buffer for 300Os. Reference sensors subjected to identical procedures without BCMA-Fc loading were included for background subtraction. Each condition was analyzed in technical triplicate to ensure data reproducibility. Raw sonograms were reference-subtracted, aligned, and globally fitted using the Gator Bio Analysis Software with a 1:1 binding model to derive the association rate constant (Kon), dissociation rate constant (Koff), and equilibrium dissociation constant (KD).

### In vitro cytotoxicity assays at multiple E:T ratios

Luciferase expressing MM.1S and MM.1R target cells were expanded, and plated in white, flat bottom 96-well plates at 1.0 × 10^4^ cells per well in 100 µL complete RPMI with 10% FBS. After an overnight settling period in incubator at 37 °C, 5% CO2 and 95% humidity. Next day, CAR T cells were added to achieve effector-to-target (E: T) ratios of 0.6:1, 1.25:1, 2.5:1 and 5:1 (otherwise specified in figures) in a final volume of 200 µL per well, each condition was performed in technical triplicate. Where indicated, synthetic peptide (nxP or P5) or conditioned “secretory-soup” from CAR cultures was added at the time of CAR T cell addition at the working concentrations used in the study. Target-only wells were included on each plate. Plates were incubated and luminescence was measured after co-culture by adding D-luciferin substrate, final concentration of 1X and reading on a Multiplate reader. Each experiment was repeated with CAR T cells derived from at least three independent donors.

### Long-term tumor rechallenge (TR) assay

A long-term TR assay was performed to evaluate the durability, persistence, and repeated antigen-stimulation capacity of CAR T cells under continuous BCMA-dependent tumor exposure, as described previously and adapted for multiple myeloma models ^42^.MM.1S target cells were rested overnight in complete RPMI-1640 medium supplemented with 10% fetal bovine serum at 37°C and 5% CO₂ to ensure recovery and exponential growth. On the following day (designated Day 0 of rechallenge), CAR T cells were co-cultured with MM.1S cells at defined E:T ratios. Initial co-cultures were established in complete AIM-V medium supplemented with IL-7 and IL-15 (84 IU and 100 IU respectively). Co-cultures were incubated at 37°C, after which cells were harvested by centrifugation (300 × g, 5 min). Culture supernatants were carefully collected and stored at -80 °C for subsequent cytokine profiling . The remaining cell pellets were resuspended and used for multiparametric flow-cytometric analysis. At each rechallenge point, CAR T cells were assessed for viability, CAR expression, exhaustion, and phenotype. Viability was determined using 7-AAD. Exhaustion markers were analysed using fluorophore-conjugated antibodies against PD-1, TIM-3, and LAG-3. Where applicable, CAR expression was assessed. Fluorescence-minus-one (FMO) controls were generated for each exhaustion marker, and all gates were stringently defined based on FMOs using FlowJo software (v10.10.0).Following analysis, viable CAR T cells were counted and fresh, exponentially growing MM.1S cells were reintroduced at the original E:T ratio to initiate the next round of rechallenge. This cycle of tumor stimulation, harvest, analysis, and reseeding was repeated sequentially throughout the duration of the assay. TR were performed on Days 9, 11, 13, 15, 17, and 19, consistent with the experimental timeline shown. Identical cell densities, media conditions, and incubation parameters were maintained across all rechallenge rounds to ensure comparability between conditions. At each rechallenge time point, samples were collected for longitudinal assessment of CAR T cell viability, expansion, exhaustion-marker expression, and functional persistence. This TR assay enabled quantitative comparison of sustained tumor control and memory-associated fitness of SHEDLOCK-engineered CAR T cells under prolonged antigenic pressure.

### Metabolic profiling

Metabolic profiling of CAR T cells was performed using the Agilent Seahorse XF T Cell Metabolic Profiling workflow on a XFe24 analyser (Agilent Technologies, Santa Clara, CA, USA) according to the manufacturer’s specifications. One day prior to the assay, sensor cartridges were hydrated overnight in a non-CO_2_ incubator in Seahorse XF Calibrant and XF PDL cell culture microplates were coated with poly-L-Lysine according to the manufacturer’s instructions. CAR T cells from running experiments were counted, washed, and co-culture experiment was set up. After the co-culture time points, the cells were collected and proceeded with CD4 and CD8 bead-based isolation and counted . The CAR T cells were then resuspended in pre-warmed Seahorse XF RPMI assay medium pH 7.4, supplemented immediately before use with 1 mM sodium pyruvate, 2 mM glutamine and 10 mM glucose (final assay medium composition per Agilent recommendations). Optimal seeding density for primary human T cells (1 lakh cells/well) was followed to achieve robust basal OCR while avoiding signal saturation. Cells were plated in XF PDL plates and allowed to settle in a non-CO_2_ incubator at 37°C for 30 mins prior to assay start. Compound stocks supplied with the kit (oligomycin, FCCP and rotenone/antimycin A) were reconstituted and diluted into assay medium following the kit instructions .Working concentrations were prepared and loaded into the hydrated sensor cartridge ports in the recommended injection order (port A: oligomycin, port B: FCCP, port C: rotenone/antimycin A) to obtain basal respiration, ATP-linked respiration, maximal respiration and non-mitochondrial respiration parameters as defined by the Agilent user guide. For glycolytic assessment, parallel wells were prepared for the Glycolysis Stress Test workflow (glucose, oligomycin and 2-DG injections) using XF RPMI medium and kit reagents, following the Agilent protocol for glucose starvation and stepwise substrate addition. All assay templates, mix/wait/measure cycle settings and plate maps were implemented using the Wave (or XF Pro Controller) templates provided with the XF T Cell kit. Real-time OCR and ECAR traces were baseline-corrected, quality-checked for outliers, and analysed using Agilent Wave or Seahorse Analytics to extract basal OCR, ATP production, maximal respiration, spare respiratory capacity, basal glycolysis, glycolytic capacity and glycolytic reserve. Data were normalized and three biological replicates from independent donor T cell preparations (n ≥ 3) were assayed in technical replicates.

### Mitochondrial ROS and membrane potential measurement

Mitochondrial ROS levels and mitochondrial membrane potential in CAR T cells were assessed by fluorescence imaging using Mito SOX Red and TMRE dye at the experimental time points indicated in the figure legends. The protocol was followed as described by us previously ^42^. Briefly, fluorescence images were acquired using EVOS imaging System (Thermo Fischer Scientific) . All staining and imaging procedures were performed under identical conditions for all groups to allow direct visual comparison between CAR T cell conditions, consistent with the representative images presented in the manuscript.

### Telomere length and telomerase activity assays

Telomere length in CAR T cells was measured by quantitative PCR using a commercial telomere length quantification kit, following the manufacturer’s instructions. CAR T cells were collected at the indicated time points after prolonged antigen exposure and in vitro TR assays. Genomic DNA was isolated, quantified, and normalized to equal input across samples. Telomeric repeats and a single-copy reference gene were amplified in parallel using SYBR Green chemistry, with reactions performed in technical triplicates. Relative telomere length was calculated as the telomere-to-single-copy gene (T/S) ratio using the ΔCt method and normalized to the corresponding control CAR T cell condition.

Telomerase activity was assessed using a real-time PCR-based TRAP assay (TRAPeze RT Telomerase Detection Kit). CAR T cells were harvested at the indicated time points, washed, and lysed in CHAPS buffer to preserve enzymatic activity. Equal amounts of protein lysate were used per reaction. Telomerase extension and amplification were performed according to the manufacturer’s protocol, with TSR8 standards included to generate a standard curve, along with appropriate negative and heat-inactivated controls. Fluorescence was acquired on a Bio-Rad CFX Opus 96 system, and telomerase activity was calculated as relative telomerase units (RTU) by interpolation from the TSR8 standard curve and normalized to control CAR T cells.

### p21 expression and senescence marker analysis

Senescence-associated p21 (CDKN1A) expression in CAR T cells was assessed by immunofluorescence staining followed by quantitative image-based analysis. CAR T cells were harvested at the indicated time points, plated on poly-L-lysine coated glass coverslips. Cells were fixed with 4% paraformaldehyde and later on permeabilized with 0.1% Triton X-100, and blocked with donkey serum. incubated with a rabbit monoclonal anti-p21 antibody (1:200), followed by an Alexa Fluor 488–conjugated anti-rabbit secondary antibody (1:500). Nuclei were mounted with antifade reagent, and images were acquired using a fluorescence microscope. p21 expression was quantified using ImageJ/Fiji by measuring nuclear integrated fluorescence intensity normalized to nuclear area. Data are reported as integrated fluorescence intensity per cell.

### Notch signaling reporter assay

Notch signaling activity was evaluated using a fluorescence-based synthetic Notch (SynNotch) reporter system^76^.Synthetic gene fragments encoding the BCMA receptor were synthesized and cloned into a pLenti expression vector, followed by lentiviral transduction into GFP-expressing HEK293 T cells to generate sender cells. A stable BCMA-expressing clone (BCMA-R-HEK293T-GFP) was isolated by fluorescence-activated cell sorting (FACS; BD Aria). For receiver cell generation, a synthetic fragment encoding a BCMA-specific single-chain variable fragment (BCMA-scFv) was cloned into the SynNotch backbone (Plasmid #79125), and lentiviral particles were used to transduce HEK293 T cells. In parallel, a reporter construct encoding pHR_Gal4UAS_tBFP_PGK_mCherry (Plasmid #79130), which enables Gal4-dependent inducible BFP expression along with constitutive mCherry expression, was introduced. Stable receiver HEK293 T cells co-expressing both constructs were isolated by FACS (BD Aria). Notch pathway activation was quantified by calculating the ratio of BFP to mCherry fluorescence intensity, with reduced BFP signal reflecting inhibition of γ-secretase-mediated Notch receptor cleavage and downstream transcriptional activation. Flow cytometry data were acquired using identical instrument settings across all experimental conditions and analyzed using FlowJo software.

### Mass spectrometry

Secreted nxP peptide in CAR T cell supernatants was confirmed by mass spectrometry, adapted from a previously described workflow^77^. CAR T cells with and without peptide (SHEDLOCK-2 and NTP control, respectively) were expanded, counted, and cultured for 24 h at a seeding density of 1 × 10⁶ CAR T cells/mL in AIM-V medium supplemented with IL-7 and IL-15 (84 IU and 100 IU respectively). After incubation, cells were centrifuged at 300 rcf for 10 min, and cell pellets and cell-free supernatants were separated to assess peptide secretion in the culture supernatant. Samples from both peptide-expressing CAR T cells and control CAR T cells were processed for mass spectrometry analysis.

MS/MS profiling was performed using an Agilent G6550B iFunnel Q-TOF LC-MS system coupled to an Agilent 1260 Infinity HPLC system. Samples (80 μL) were precipitated with 1 mL TCA–acetone and incubated at 4 °C for 2 h. Protein pellets were dissolved in lysis buffer, and 80 μg of protein from each sample was resolved on a 15% SDS-polyacrylamide gel with a 4% stacking gel using a vertical electrophoresis apparatus at 20 °C under constant current (50 mA/gel) until the bromophenol blue dye front reached the bottom of the gel. Gels were visualized by Coomassie Brilliant Blue (CBB) staining. Excised gel bands were reduced with 5 mM TCEP, alkylated with 50 mM iodoacetamide, and digested with trypsin (1:50 trypsin-to-lysate ratio) for 16 h at 37 °C. Digested peptides were desalted using a C18 silica cartridge and dried using a SpeedVac. The dried pellets were resuspended in buffer (5% acetonitrile, 0.1% formic acid).

Samples were centrifuged at 8, 000 rpm for 5 min, and the clarified supernatant was transferred to LC vials. For LC-MS/MS analysis, 1 μg of peptide digest was injected onto an Agilent AdvanceBio Peptide Mapping C18 column (2.1 × 150 mm, 2.7 μm particle size) maintained at 40 °C. Peptides were eluted using a 0–40% gradient of buffer B (80% acetonitrile, 0.1% formic acid) at a flow rate of 300 μL/min over a 60-min gradient. The G6550B Q-TOF was operated in positive-ion Auto MS/MS mode with the following parameters: capillary voltage 3, 500 V; fragmentor voltage 175 V; acquisition rate 3 spectra/s (MS) and 3 spectra/s (MS/MS); precursor selection by abundance with a maximum of 5 precursors per cycle; dynamic exclusion enabled for 10 s after 2 spectra; and collision energy applied using a slope/offset formula optimised for tryptic peptides. Data acquisition and analysis were performed using the Agilent MassHunter Qualitative Analysis and BioConfirm suite. Detected peptides are listed in detail in Table 6.

### RNA sequencing and pathway enrichment (GSEA)

RNA sequencing was performed on CAR T cells harvested at the indicated experimental endpoints to define transcriptional programs associated with SHEDLOCK-2 versus NTP-CAR phenotypes. Total RNA was isolated using the QIAGEN RNeasy Mini kit with on-column DNase treatment and mRNA was enriched using QIAGEN Pure mRNA beads; strand-specific libraries were prepared following heat-fragmentation and adapter ligation and libraries were purified for Illumina NovaSeq 6000 compatibility as described. Libraries were sequenced on an Illumina NovaSeq platform and raw fasta data were processed through the lab’s standard nf-core preprocessing workflow; differential expression and normalization were performed using DESeq2 package in R studio. Ranked gene lists derived from the differential analysis were subjected to pathway enrichment and gene-set testing, and the principal pathways highlighted in the figures (glycolysis, oxidative phosphorylation and FoxO signaling) were identified by GSEA and corroborated by complementary enrichment/annotation tools (DAVID, KEGG and ShinyGO) as reported in Supplementary Figure S20 and **Table 7**. Resulting heatmaps, PCA plots and enrichment plots were generated using the lab’s standard visualization workflow and interpreted using normalized enrichment scores and the adjusted significance thresholds employed in the manuscript.

### In vivo xenograft mouse model

To establish systemic multiple myeloma xenografts, mice were injected intravenously via the tail vein on day 0 with luciferase-expressing MM.1S cells at the indicated cell numbers, as specified in the corresponding figure legends. Tumor engraftment and progression were monitored longitudinally by bioluminescence imaging (BLI) at predefined time points. For in vivo validation of the γ-secretase-inhibitory peptide P5, NSG mice bearing MM.1S tumors received a single intravenous administration of P5 on day 12 post-tumor inoculation. P5 was delivered at doses of 1 mg, 2 mg, or 4 mg per mouse, formulated in sterile phosphate-buffered saline (PBS). Control mice received vehicle alone. On day 15, mice were euthanized, and blood was collected by cardiac puncture. Bone marrow was harvested by flushing femurs and tibias with sterile buffer to generate single-cell suspensions. Plasma isolated from peripheral blood and cell-free supernatants obtained from processed bone marrow samples were used for quantification of soluble BCMA (sBCMA) levels by ELISA. In parallel, isolated myeloma cells were stained with fluorophore-conjugated anti-BCMA (CD269) antibody and 7-AAD to assess membrane BCMA (mBCMA) expression and cell viability by flow cytometry.

For evaluation of in vivo anti-tumor efficacy and CAR T cell persistence, MM.1S-bearing mice received NTP-CAR T cells, SHEDLOCK-1 CAR T cells, or SHEDLOCK-2 CAR T cells at the indicated doses and time points, as defined in the experimental schematics. Tumor burden was monitored serially by BLI, while survival was recorded over the duration of the study. Peripheral blood and bone marrow were collected at defined intervals to quantify circulating and tissue-resident CAR T cells by flow cytometry and to assess CAR expression, persistence, and immunophenotype.

To assess functional durability under repeated antigen exposure, a tumor-rechallenge (TR) model was employed. On day 0, mice were engrafted with 3 × 10^5^ MM.1S cells and on day 10 they received a single infusion of 1 × 10^6^ patient-derived CAR T cells. Following initial tumor clearance after primary CAR T cell infusion, mice were re-challenged intravenously with 0.1 × 10^6^ luciferase-expressing MM.1S cells on day 15 post-CAR T infusion, as depicted in the study timeline. No additional CAR T cells were administered at the time of rechallenge. Tumor progression was monitored longitudinally by bioluminescence imaging (BLI). At defined endpoints following rechallenge, single-cell suspensions were prepared from peripheral blood, spleen, and bone marrow for comprehensive phenotypic and functional profiling of CAR T cells, including assessment of exhaustion markers (PD-1, TIM-3, and LAG-3) and phenotypic status into naïve, effector, effector-memory, and central-memory subsets based on CD45RO and CCR7 expression.

Another TR model was established in NCG mice to track CAR T cell metabolic and telomere parameters against therapeutic outcome. Tumor re-challenge was performed by repeated injection of 1 × 10^5^ MM.1S cells on days 15, 20, and 25, and animals were followed for survival. Two complementary cohorts were used. In the longitudinal mechanistic cohort, CAR T cells generated from five individual patient samples (P1 to P5) were tested with 6 NCG mice per group for each patient sample, for a total of 30 mice, and cells were sampled before infusion (day 10) and at serial post-administration time points (days 12, 16, 21, and 26) for kinetic analysis of each parameter. In the extended correlation cohort, CAR T cells were generated from 15 patient samples and, where indicated, engineered as SHEDLOCK-2 cells, with 4 mice infused per group and pooled across patients for a total of 60 NCG mice. Pre-administration measurements (day 10) included telomere length, telomerase activity, Hes1 expression, and mitochondrial OCR and ECAR, while post-administration measurements captured survival, CAR T cell persistence, and CAR T cell exhaustion. At each time point, blood and bone marrow were collected and human CD8⁺ T cells were isolated by magnetic bead selection. Recovered cells were co-cultured with MM.1S target cells for 24 h, after which cells and supernatants were harvested for downstream analysis.

### Correlation analysis

To test whether these parameters predict outcome, pre-infusion and post-TR values for telomere length, telomerase activity, OCR, ECAR, Hes1 expression, PD-1, CAR T persistence, and survival were compiled across the 15-patient cohort. Pairwise relationships were evaluated by Pearson correlation, and the full set of comparisons was summarized as a bubble correlation matrix, with representative pairwise regressions shown alongside the correlation coefficient (r) and p value for each comparison. Correlations with p < 0.05 were considered significant.

### Serial transfer CAR T cell mouse model

A serial-transfer mouse model was developed to evaluate the long-term persistence of SHEDLOCK-engineered CAR T cells, as schematically illustrated in *Figure 7*. NSG mice bearing established MM.1S xenografts and treated with primary CAR T cell infusion were humanely euthanized on day 10 post CAR T cell administration, corresponding to the peak persistence phase, and CAR T cells were enriched using CD3 magnetic bead-based selection and designated as serially transferred CAR T cells (CAR T-ST). A total of 0.5 X 10^6^ CAR T-ST cells were intravenously infused into secondary NSG mice that had been pre-engrafted 10 days earlier with 0.3X10^6^ MM.1S cells. Tumor control, CAR T cell persistence, and survival were evaluated longitudinally across NTP, SHEDLOCK-1 and SHEDLOCK-2 CAR T cells.

### Patient-derived xenograft mice model

A patient-derived multiple myeloma model was established in busulfan-conditioned NSG-exIL6 mice to evaluate CAR T cell efficacy in a clinically relevant setting, following the published protocol^45^ . Briefly, 6-8-week-old NSG mice were conditioned with busulfan (30 mg/kg, intraperitoneally) and maintained on human IL-6 supplementation (recombinant IL-6, 10 µg per mouse, single intraperitoneal dose) to support plasma-cell survival. Primary myeloma cells obtained from patient bone-marrow aspirates were processed immediately after collection, and malignant plasma cells were enriched using the RosetteSep Human Multiple Myeloma Cell Enrichment Cocktail (STEMCELL Technologies), as described above. The enriched CD38⁺/CD138⁺ myeloma cell fraction was counted, assessed for viability, and 1 × 10⁶ cells per mouse were injected intravenously in sterile PBS on day 0.Tumor engraftment and disease progression were monitored longitudinally by measurement of circulating human immunoglobulin levels in plasma and by flow-cytometric detection of human CD38⁺/CD138⁺ cells in peripheral blood, with progressive disease typically evident by week 5-6 post-engraftment. Therapeutic interventions, including CAR T cell infusion, were performed at the time points specified in the figures and figure legends..

### Histopathological evaluation of major organs

At study termination, mice were euthanized according to institutional guidelines and a complete necropsy was performed. Major organs, liver, kidney, lung, heart, spleen, brain, and both femora, were harvested, rinsed briefly in cold PBS to remove blood, and immersion-fixed in 10% neutral-buffered formalin, as described by us previously^78^.Histologic evaluation scored parameters including inflammatory cell infiltration, necrosis, tissue architecture disruption, haemorrhage, and tumor infiltration on a semi-quantitative scale (0 = absent, 1 = minimal, 2 = mild, 3 = moderate, 4 = severe). For bone sections, marrow cellularity and tumor burden were assessed and reported as percent infiltration.

### Clinical chemistry for hepatic and renal function

Blood samples were collected from xenografted or treatment-group mice at defined experimental endpoints via retro-orbital or terminal cardiac puncture into serum-separation tubes. Samples were allowed to clot at room temperature for 20-30 minutes and centrifuged at 2, 000 × g for 10 minutes to obtain serum. Serum biochemical parameters were measured using a fully automated clinical chemistry analyzer in accordance with the manufacturer’s guidelines. Hepatic function was assessed by quantifying alanine aminotransferase (ALT), aspartate aminotransferase (AST), and alkaline phosphatase (ALP). Renal function was evaluated by measuring serum creatinine, blood urea nitrogen (BUN), and uric acid. All assays were performed in technical duplicates and included internal laboratory controls to ensure measurement accuracy. Data were analyzed relative to treatment cohorts to identify any peptide- or CAR T cell-associated toxicity, and results were interpreted in conjunction with histopathology and cytokine-release measurements to provide a comprehensive assessment of systemic safety.

### Cytokine release syndrome (CRS) mice model

Cytokine release syndrome (CRS) was modelled in MM.1S tumor–bearing immunodeficient mice following CAR T cell infusion. Mice bearing established MM.1S tumors (0.5 × 10⁶ cells, intravenous) received a single intravenous infusion of 5 × 10⁶ CAR T cells per mouse (NTP-CAR T cells or SHEDLOCK-1 CAR T cells) on day 10 post-tumor engraftment, as depicted in Supplementary Figure S11. To evaluate IL-6 pathway blockade, a subset of SHEDLOCK-1–treated mice received tocilizumab (anti–IL-6 receptor antibody) administered as a single intraperitoneal dose of 10 µg per mouse at the time of CAR T cell infusion. Following CAR T cell administration, mice were monitored for clinical signs associated with CRS. Peripheral blood was collected on day 15, and plasma was isolated for downstream analysis. Serum cytokine levels, including IL-1β, TNF-α, IFN-γ, IL-2, IL-6, and GM-CSF, were quantified to assess CRS-associated inflammatory responses. In parallel, serum clinical chemistry parameters (ALT, AST, ALP, and albumin) were analysed to evaluate systemic toxicity.

### Sub-chronic (90-day) safety assessment

To evaluate long-term safety, a sub-chronic study was performed over 90 days in NCG mice. SHEDLOCK-2 or NTP-CAR T cells were tested, alongside a sham control group that received vehicle (un-transduced T cells). Tumor-bearing mice were infused with SHEDLOCK-2 CAR T cells and monitored for the full study period, with blood and tissues collected at the terminal time point. Serum cytokines were measured by multiplex immunoassay or ELISA and included IL-10, IL-17A, IL-12p70, and MCP-1 in addition to the original panel of IL-6, TNF-α, IL-1β, IL-2, GM-CSF, and IFN-γ. Complete blood counts were obtained on an automated hematology analyzer to assess hematopoiesis. Organ function was evaluated from serum using cardiac troponin-I as a sensitive marker of cardiac injury, cystatin-C together with creatinine and blood urea nitrogen (BUN) for renal function, and ALT, AST, ALP, and albumin for hepatic function. At necropsy, liver, kidney, lung, heart, spleen, intestine, brain, and bone marrow were fixed in 10% neutral-buffered formalin, embedded in paraffin, sectioned, and stained with hematoxylin and eosin. A blinded, board-certified pathologist reviewed all sections and assigned semi-quantitative lesion scores on a 0 to 4 scale, with values compared against sham controls to identify any drug-related findings.

### Image illustrations

All schematic illustrations presented in this study were created either wholly or in part using Bio Render. Created in Bio Render. Chauhan, V. (2026) https://biorender.com/ty7zpv

## Data Availability

All data generated or analyzed in this study, including unique reagents, are available from the authors upon reasonable request. To protect patient confidentiality, individual clinical data are not publicly released; however, de-identified participant-level data may be obtained from the corresponding author upon request. RNA-sequencing and mass spectrometry datasets have been deposited in a public repository (Zenodo link: 10.5281/zenodo.21883517).

## Code Availability

All computational scripts and AI-driven modeling pipelines used in this work have been deposited in a public GitHub repository. Supplementary code supporting data analysis and figure generation is available from the authors upon reasonable request.

## Statistical analysis

All analyses were performed in GraphPad Prism v10.3. Normality was assessed by Shapiro-Wilk; as most datasets deviated from normality, non-parametric tests were used unless stated otherwise in the figure legend. Two groups were compared by two-tailed Mann-Whitney U test and three or more by Kruskal–Wallis with Dunn’s correction. Two-factor experiments, including effector-to-target ratios and longitudinal tumor-rechallenge data, were analysed by two-way ANOVA with Šídák’s correction. Survival was compared by log-rank (Mantel-Cox) test on time-to-endpoint data, with mice alive at study end censored at last follow-up, and is shown per mouse. Correlations used Pearson on donor-level means, the only parametric exception. Individual tumor-burden trajectories are shown descriptively, without formal comparison.

Data are presented as mean ± SEM, with *P < 0.05, **P < 0.01, ***P < 0.001 and ****P < 0.0001. Throughout the manuscript, n denotes independent biological replicates, that is separate healthy donors, patient samples or mice, and not technical replicates, with each condition set up in technical duplicate or triplicate within an experiment. All in vitro assays used cells from at least three independent donors and were repeated at least three times, and the number of patient samples and of mice per group is stated in the corresponding figure legend.

## Acknowledgement

The authors gratefully acknowledge the contributions of all TA Lab members: Dr. Areej Akhtar, Md Shakir, Mehak Pracha, Aashi Singh, Mohd Nadeem, Prial Taneja, Md Imam Faizan, Iqbal Azmi and Ubaid Akhtar. We also thank Dr. Garima Nirmal for her assistance in procuring human samples. We sincerely acknowledge the staff of CSIR-Indian Institute of Chemical Biology (CSIR-IICB) and MicroCRISPR Pvt. Ltd. for their critical support in the pre-clinical studies and also Cellogen therapeutics for providing the funding. We thank National Institute of Immunology for the Seahorse assay under grant no. NII-471 (EQPT 27171), supervised by Dr. Aneesh Kumar AG at the Central Instrumentation Facility. We thank Optical Microscopy Facility of the Advanced Technology Platform Centre (ATPC), Regional Centre for Biotechnology, supported by the Department of Biotechnology (DBT) (Grant No. BT.MED-II/ATPC/BSC/01/2010), for Super-resolution microscopy under the supervision of Mr. Suraj Tewari and Ms. Damini. A patent application related to this work has been filed by TA and SS. We also acknowledge the use of Claude.ai for grammar and language refinement during manuscript preparation; all scientific content, analyses, and interpretations were conceived, generated, and validated by the authors.

## Author Contributions Statement

T.A. conceived and supervised the study. S.S. performed the majority of the experiments, while V.C. and Z.B. conducted all computational analyses. J.A. and A.K.S. performed and supervised the in vivo studies. D.M. contributed to mass spectrometry and transcriptomic data analysis. S.G., D.R.S., R.A., M.S.A., D.D., P.P., N.C., D.K.S., and V.P.S. assisted with experimental execution, biolayer interferometry, computational analyses, and data interpretation. M.H., U.M., and S.P.R., provided technical expertise and contributed to manuscript editing. G.K., and D.P., facilitated access to clinical samples and provided clinical oversight. T.A. and A.K.V. supported study interpretation and overall project guidance. S.S. and T.A. wrote the manuscript, and all authors reviewed and approved the final version.

